# CDK8 phosphorylation of DELLA limits Mediator recruitment in gibberellin signaling

**DOI:** 10.64898/2026.05.31.729123

**Authors:** Xu Huang, Hongyi Chen, Larry Reser, Andres V. Reyes, Shou-Ling Xu, Jeffrey Shabanowitz, Donald F. Hunt, Tai-ping Sun

## Abstract

Gibberellin (GA) promotes plant growth primarily by triggering degradation of DELLA transcription regulators, yet how DELLA activity is fine-tuned dynamically by phosphorylation independently of proteolysis remains poorly understood. Here we show that the CDK8 kinase module of the Mediator complex attenuates activity of the *Arabidopsis* DELLA protein RGA by modulating coactivator recruitment. Using TurboID-based proximity labeling, biochemical and genetic analyses, we identify CDK8 as an in-planta kinase that phosphorylates RGA at Ser170 within its disordered PolyS/T region. This phosphorylation does not affect RGA stability, localization or interactions with transcription factors or histone H2A, but selectively weakens RGA association with the Mediator subunit MED15, thereby reducing DELLA-dependent transcription activation. Consistently, *cdk8* mutants show impaired GA-responses and delayed developmental phase transitions that are partially rescued by loss of DELLA function. Our findings uncover a phosphorylation-dependent mechanism by which the Mediator kinase module fine-tunes hormone-responsive transcription through selective control of DELLA–coactivator interactions.

## INTRODUCTION

The phytohormone gibberellin (GA) is a central regulator of plant development, controlling seed germination, vegetative growth, floral transition, and reproductive development^1^. GA perception is mediated by the receptor GIBBERELLIN INSENSITIVE1 (GID1), which triggers the ubiquitin–proteasome-dependent degradation of DELLA proteins, the principal repressors of GA signaling^2,3^. Dominant mutations in *DELLA* genes underlie the semi-dwarf, high-yielding wheat varieties that fueled the Green Revolution of the 1960s, highlighting the key role of DELLA proteins in regulating plant growth^4,5^. First identified in *Arabidopsis thaliana* as repressor of GA signaling^6,7^, DELLAs are evolutionarily conserved across land plants^8^ and function as integrative hubs that coordinate hormonal and environmental signals to balance growth, development, and stress responses^5,9^.

DELLA proteins are plant-specific GRAS family of transcription regulators, characterized by a conserved C-terminal GRAS domain^6,7,10^ and a unique N-terminal DELLA domain required for GA-dependent proteolysis^11,12^. Upon GA binding, GID1 forms a ternary GA–GID1–DELLA complex that recruits the SCF^SLY1/GID2^ E3 ubiquitin ligase, leading to DELLA polyubiquitination and rapid degradation, and thereby enabling transcriptional reprogramming of GA-responsive genes^2,13–19^. While this proteolytic pathway is a major determinant of DELLA abundance, accumulating evidence indicates that DELLA activity can also be modulated independently of protein degradation^5,18^.

At the molecular level, DELLA proteins regulate gene expression primarily through protein–protein interactions rather than direct DNA binding. Through their GRAS domain, they associate with a wide array of transcription factors (TFs) and chromatin remodelers to reprogram transcriptional outputs^9,20–23^. Recent work has shown that DELLA function requires the formation of TF–DELLA–histone H2A complexes at target loci, enabling both transcriptional repression and activation depending on the interacting partners^24^. Notably, DELLA-mediated transcriptional activation depends on recruitment of the Mediator coactivator complex via interaction with the tail subunit MED15, a mechanism conserved across land plants^25^. How DELLA interactions with Mediator and other cofactors are dynamically regulated remains largely unknown.

Structurally, the GRAS domain comprises five conserved subdomains—Leucine Heptad Repeat 1 (LHR1), VHIID, Leucine Heptad Repeat 2 (LHR2), PFYRE, and SAW—three of which are named after conserved sequence motifs^10^. Functional and structural studies have defined distinct roles to these subdomains: LHR1 mediates interactions with TFs to recruit DELLA proteins to target promoters, whereas the PFYRE subdomain stabilizes chromatin-associated complexes through interaction with histone H2A^9,24^. In contrast, the VHIID, LHR2 and SAW subdomains interact with F-box proteins, linking these regions to DELLA turnover via the ubiquitin–proteasome pathway^18,26–28^. Together, these findings highlight the modular architecture of DELLA proteins and suggest multiple layers of regulatory control.

In addition to GA-induced degradation via polyubiquitination, DELLA activity is regulated by multiple post-translational modifications (PTMs), including Small Ubiquitin-Like Modifier-conjugation (SUMOylation), glycosylation, and phosphorylation^29–32^. Among these, phosphorylation has been proposed as an important regulatory layer, although its functional role remains unclear^5,29^. Early studies suggested that phosphorylation of the rice DELLA SLR1 promotes interaction with the F-box protein GID2 and thereby facilitates proteasomal degradation^14,33,34^, but later work found no effect of phosphorylation on GID2 binding, arguing against a direct role in DELLA turnover^35^. Consistent with this view, pharmacological inhibition of phosphatases or genetic suppression of the protein phosphatase TOPP4 attenuated GA-induced DELLA degradation, suggesting that dephosphorylation rather than phosphorylation may favor proteolysis^36–39^. Several kinases, including casein kinase I EARLIER FLOWERING1 (EL1) and a GSK3/SHAGGY-like kinase, have been reported to phosphorylate DELLA proteins and enhance their stability, although these activities have only been demonstrated in vitro^40,41^. More recently, mass spectrometry analyses of *Arabidopsis* DELLA, RGA (for REPRESSOR OF *ga1-3*), have identified multiple phosphorylation sites within the intrinsically disordered PolyS and PolyS/T regions flanking the DELLA domain, and mutational studies showed that phosphorylation in these regions modulates DELLA activity without affecting protein stability by promoting association with histone H2A at target loci^42^. However, the identity of the protein kinase(s) responsible for DELLA phosphorylation in planta, and the mechanistic consequences of specific phosphorylation events, remain largely unknown^5^.

Here, using TurboID-based proximity labeling^43^ together with biochemical and genetic approaches, we identify the CDK8 kinase module (CKM) of the Mediator complex as an in-planta regulator of DELLA phosphorylation and uncover a broad set of DELLA-associated transcription regulators. We show that CDK8 phosphorylates the *Arabidopsis* DELLA protein RGA at Ser170 within its intrinsically disordered PolyS/T region, attenuating RGA transactivation activity by weakening its interaction with MED15. These findings uncover a mechanism by which the Mediator kinase module fine-tunes hormone-responsive transcription through phosphorylation-dependent control of DELLA–coactivator interactions.

## RESULTS

### Proximity labeling reveals the RGA proximitome

To identify kinase(s) that associate with the *Arabidopsis* DELLA protein RGA in planta, we performed TurboID (TbID) proximity labeling^43–45^ to capture target proteins such as protein kinases that may only transiently interact with RGA in planta. We generated transgenic *Arabidopsis* expressing *P_RGA_:His-FLAG-RGA-TbID* in *ga1-13 della pentuple* (*ga1 dP)* background (*RGA-TbID* line). The *ga1-13* null allele causes a GA-deficient dwarf phenotype, whereas *ga1-13 dP* plants are tall due to the removal of all five *DELLA* genes. The RGA-TbID fusion protein in *ga1 dP* restored the *ga1* dwarf phenotype and underwent GA-induced degradation similar to FLAG-RGA, indicating normal functionality (**Figures S1A-1D**). As a control for nonspecific labeling, we generated a *P_RGA_:His-FLAG-NLS*-*GFP-TbID* line in the *ga1 dQ* (quadruple *della* with WT *RGA*) background (*GFP-TbID* line) with comparable expression levels as RGA-TbID (**Figures S1A-1D**). Seedlings were treated with 100 µM biotin for 3 h (**Figure S1E)**, and biotinylated proteins were purified with streptavidin beads and identified by liquid chromatography-tandem mass spectrometry (LC-MS/MS) with label-free quantification (LFQ). Proteins significantly enriched in *RGA-TbID* relative to *GFP-TbID* were identified from three biological replicates using statistical thresholds of FDR < 0.01 and S0 = 0.8 (**Figure 1A** and **Table S1**). Among the 199 RGA-proximal proteins, Gene Ontology (GO) analysis revealed strong enrichment for nuclear factors involved in transcriptional regulation, including transcription factors and regulators, Mediator complex components, chromatin remodelers, and histone-modifying enzymes (**Figure 1B, Tables S1-S2**). The specificity of the labeling is validated by the detection of many known RGA-interacting transcription factors/regulators^3,22^, including BZR1, IDDs, EIN3, TCPs, ARFs, ARR1, bHLHs and JAZ9, as well as the Mediator tail subunit MED15^25^ and the chromatin remodelers BRAHMA (BRM)^46^ and SWI3C^47^ (**Figure 1C** and **Table S1**).

**Figure 1.**
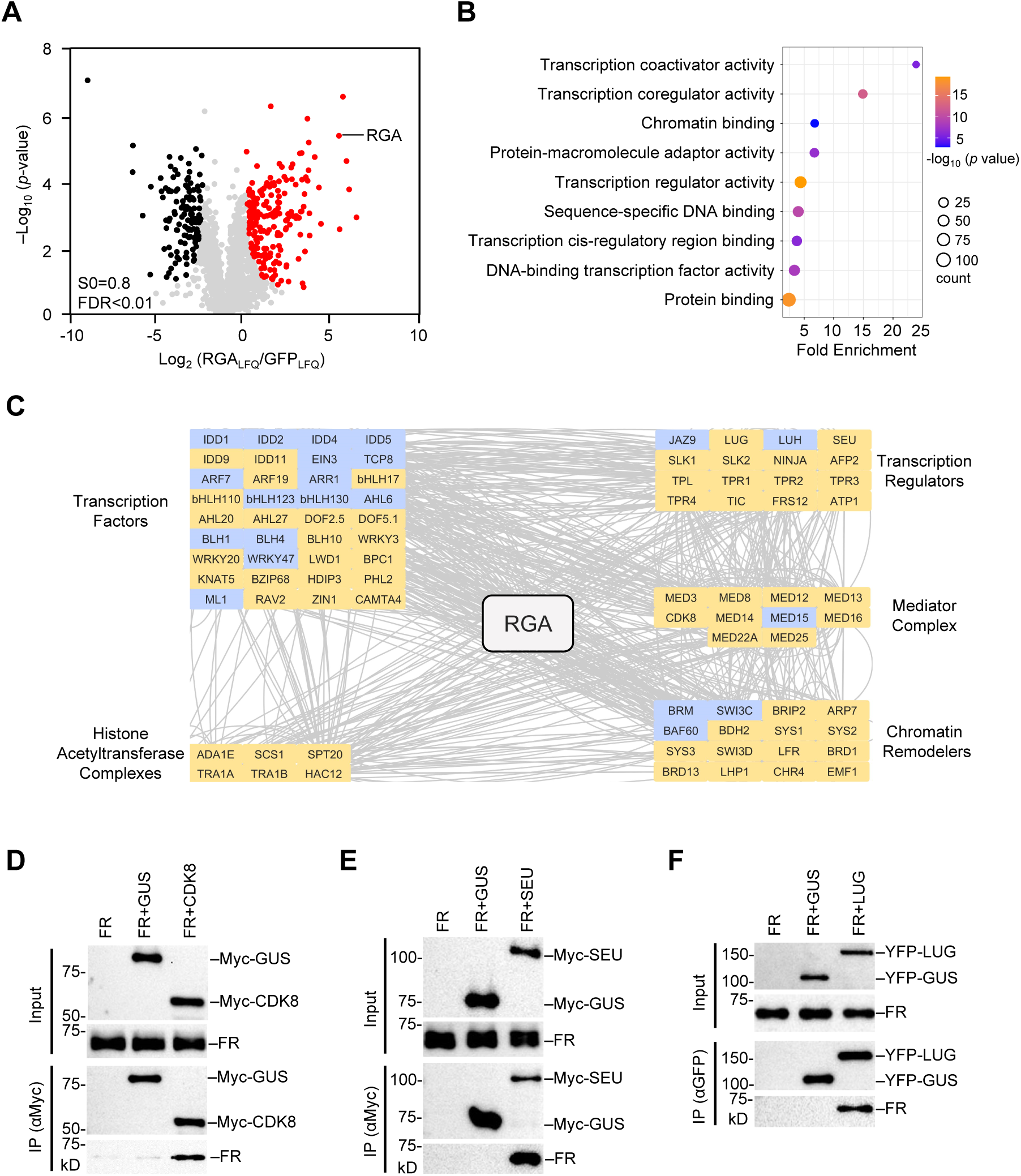
RGA proximitome identified by TbID-labeling. (A) Volcano plot of RGA proximal proteins, identified by label-free quantification (LFQ) MS analysis of biotinylated proteins purified from RGA-TbID and GFP-TbID samples. Proteins significantly enriched in RGA-TbID are labeled in red, while those enriched in GFP-TbID are in black. (B) Enriched molecular functions (MFs) of RGA proximal proteins by GO term analysis. (C) STRING (Search Tool for the Retrival of Interacting Genes/Proteins) network analysis^82^ of RGA proximal proteins involved in transcription regulation. Known RGA interactors^22^ are labeled in blue and new RGA proximal proteins are in yellow. (D)-(F) Co-IP assays showing direct interactions between RGA and three proximal proteins CDK8, SEU and LUG. FLAG-RGA was expressed alone or together with Myc-CDK8, Myc-SEU, or -YFP-LUG, separately, in *N. benthamiana*. Myc-GUS or YFP-GUS was included as a negative control. In (D-E), Myc-GUS, -CDK8 or -GUS were immunoprecipitated using anti-Myc agarose. In (F), YFP-GUS and-LUG were immunoprecipitated using anti-GFP agarose. Immunoblots containing input and IP eluates were probed with anti-FLAG, anti-Myc or anti-GFP antibodies. Representative images of 2 biological repeats are shown.

Notably, the catalytic subunit of the Mediator kinase module, CDK8, was strongly enriched together with the module components MED12 and MED13^48,49^, identifying the CDK8 kinase module (CKM) as a candidate regulator of RGA (**Figure 1C** and **Table S1**). Additional Mediator subunits, MED14 and MED16, were also identified along with MED15. The RGA proximitome further include transcriptional cofactors such as SEUSS (SEU) and SEU-LIKEs (SLKs)^50,51^ and Groucho/Tup1-type corepressors LEUNIG (LUG)/LUG HOMOLOG (LUH) and TOPLESS (TPL)/TPL-RELATEDs (TPRs)^52^, and components of histone-acetyltransferase complexes, including HAC12 in the CBF family, SAGA complex subunits (SCS1, TAF12B, and PHL/SPT20), and NuA4 complex components TRA1A and TRA1B^53,54^ (**Figure 1C** and **Table S1**). To validate these candidates, CDK8, SEU, and LUG were tested by co-immunoprecipitation (IP) assays and each showed direct interaction with RGA when co-expressed in *Nicotiana benthamiana* (**Figures 1D-1F**).

Together, these results indicate that RGA associates with transcriptional regulatory machinery comprising transcription factors, Mediator components, transcription cofactors, chromatin remodelers, and histone acetyltransferase complexes. Among these candidates, the CDK8 kinase module was of particular interest because it contains a catalytic kinase, suggesting a potential mechanism for DELLA phosphorylation.

### Phosphorylation of RGA by CDK8 kinase module

To examine whether CDK8 phosphorylates RGA in planta, we compared phosphorylation levels of endogenous RGA in WT vs *cdk8* mutants using Phos-tag SDS-PAGE because phospho-RGA (pRGA) could not be separated clearly from the unphosphorylated form by standard SDS-PAGE^42^. Phos-tag gel analysis showed that pRGA levels were reduced in both *cdk8-1* and *cdk8-2* mutants compared with WT, and RGA phosphorylation was restored in the *35S:Myc-CDK8 cdk8-1* transgenic line (**Figure 2A**). To facilitate monitoring of RGA phosphorylation, we generated transgenic *Arabidopsis* (Col-0 ecotype) carrying *P_RGA_:His-FLAG-RGA^GKG^* in *ga1-13* or *ga1-13 cdk8-1* backgrounds. His-FLAG-RGA^GKG^ was previously shown to be functional in planta^55^; RGA^GKG^ contains an additional trypsin cleavage site created by inserting a Lys (K) residue within the PolyS/T region, enabling MS detection of this region^55^. Similar to the endogenous RGA, FLAG-RGA^GKG^ showed reduced phosphorylation in the *cdk8* mutant compared with WT (**Figure 2B**).

**Figure 2.**
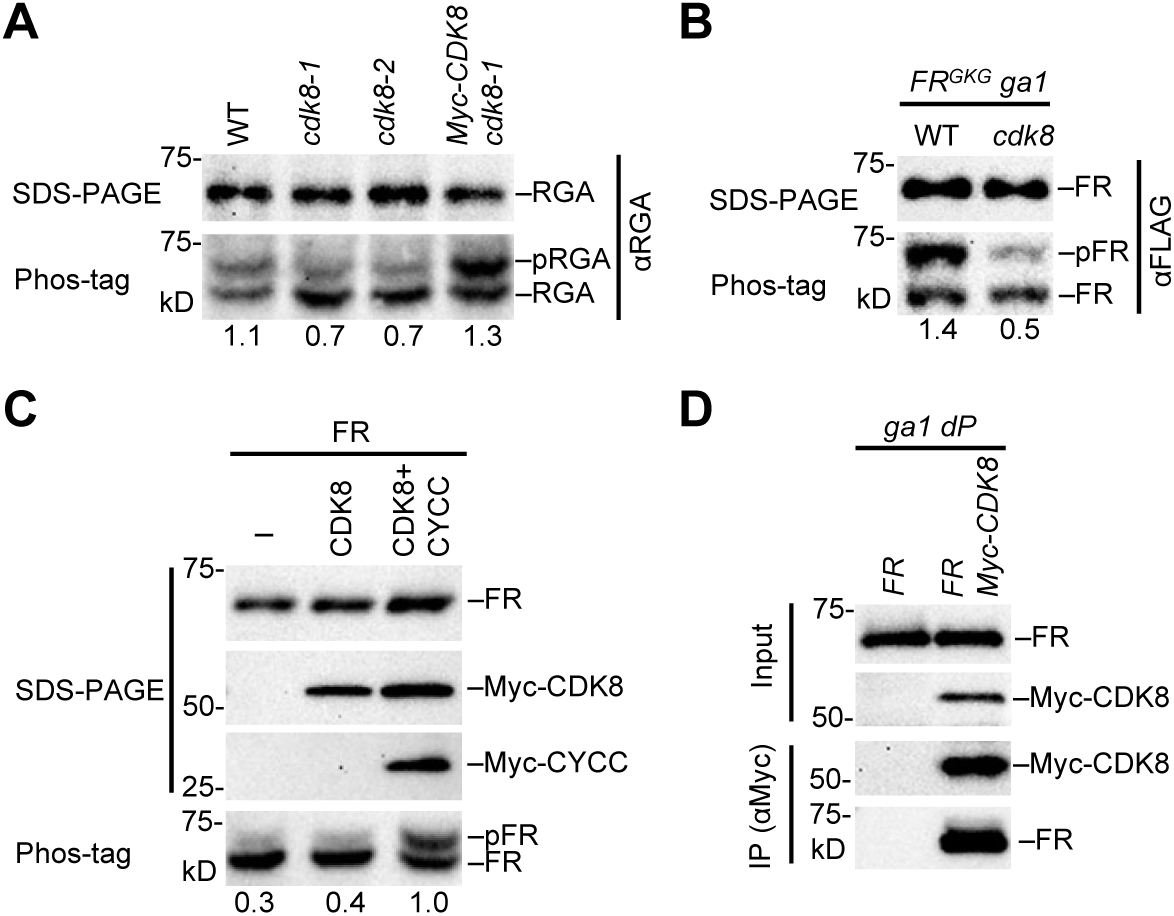
RGA Phosphorylation by CDK8 kinase module. (A) *cdk8* mutations reduced phosphorylation of endogenous RGA, while *35S:Myc-CDK8* in transgenic *Arabidopsis* rescued this mutant defect. pRGA, phosphorylated RGA. Protein blots contained total proteins extracted from 10d-old *Arabidopsis* seedlings that were grown in medium containing 1 μM paclobutrazol (PAC, a GA biosynthesis inhibitor). (B) Phosphorylation of His-FLAG-RGA^GKG^ (FR) was reduced by *cdk8*. pFR, phosphorylated His-FLAG-RGA. (C) Co-expression of both Myc-CDK8 and -CYCC increased His-FLAG-RGA (FR) phosphorylation in *N. benthamiana*. FR was expressed alone (–) or co-expressed with Myc-CDK8 or both Myc-CDK8 and -CYCC. (D) Co-IP assays showing CDK8-RGA interaction in *Arabidopsis*. Immunoprecipitation was performed using anti-Myc conjugated beads and protein extracts from transgenic lines in the *ga1 dP* background carrying *P_RGA_:His-FLAG-RGA (FR)* alone or together with *P_Ubq10_:Myc-CDK8*. Immunoblots containing input *Arabidopsis* extracts and immunoprecipitated samples were detected with anti-Myc and anti-FLAG antibodies, separately. In (A)-(C), the ratios of phosphorylated RGA/unphosphorylated RGA are shown below the Phos-tag gel blots. In (A)-(D), representative images of two biological repeats are shown.

CKM consists of CDK8, Cyclin C (CYCC), MED12 and MED13^48,49^. We found that co-expression of both CDK8 and CYCC with FLAG-RGA (FR) in *N. benthamiana* dramatically enhanced pFR levels, whereas co-expression of CDK8 alone did not (**Figure 2C**). Addition of the remaining two CKM subunits, MED12 and MED13, in the co-expression assay did not further increase pFR levels (**Figure S2**), indicating that these two CKM subunits are not limiting in this heterologous system. As described above, Myc-CDK8 interacted with FLAG-RGA when they were co-expressed in *N. benthamiana* (**Figure 1D**). Co-IP assays further show that the VHIID subdomain in RGA is required for CDK8 interaction (**Figure 3**). To detect their interaction in *Arabidopsis*, co-IP assays were performed using transgenic *Arabidopsis* in the *ga1 dP* background carrying *P_RGA_:His-FLAG-RGA* alone or together with *P_Ubq10_:Myc-CDK8* (**Figure 2D**). His-FLAG-RGA was co-immunoprecipitated with Myc-CDK8. Taken together, these results support that RGA is phosphorylated by CDK8 of the CKM.

**Figure 3.**
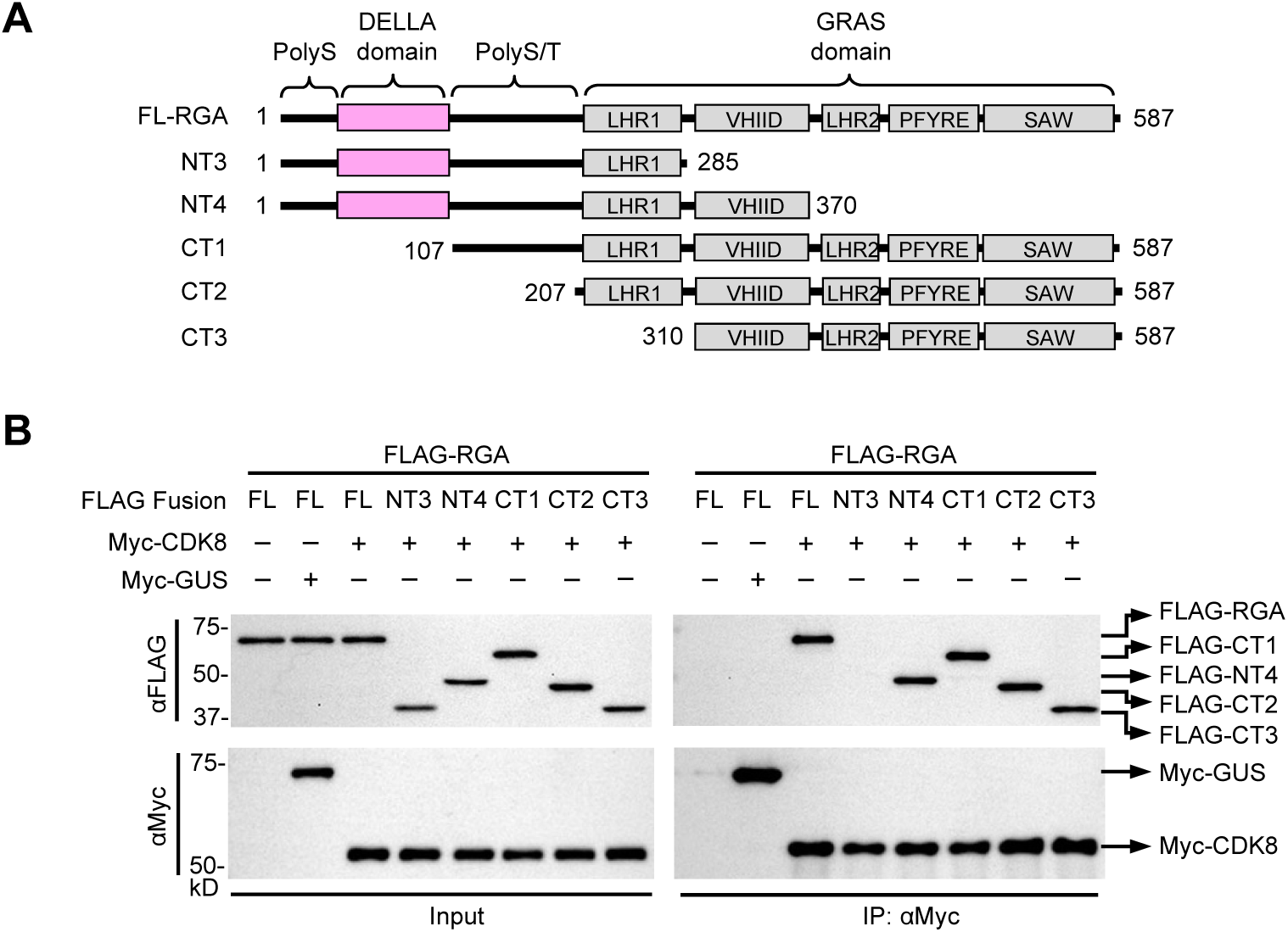
CDK8 interacted with VHIID subdomain of RGA. (A) Schematics of full-length (FL) and truncated RGA proteins. LHR, Leucin heptad repeat. (B) Co-IP assays showed that VHIID subdomain is required for CDK8 binding. FLAG-RGA (FL or truncated) was expressed alone (–) or together with Myc-CDK8 in *N. benthamiana*. Myc-GUS was included as a negative control. Myc-CDK8 or -GUS were immunoprecipitated using anti-Myc agarose. Left panel (Input): Immunoblots containing total protein extracts. Right panel (αMyc immunoprecipitated samples): Immunoblots containing IP eluates were probed with anti-FLAG or anti-Myc antibodies as labeled. Representative images of 2 biological repeats are shown.

### CDK8 promotes GA signaling by inhibiting RGA function

The role of CDK8 in regulating RGA activity and GA response in planta was analyzed by hypocotyl elongation assay. Importantly, the *cdk8* mutants displayed a reduced GA response compared with WT (**Figures 4A-4B**), suggesting that CDK8 promotes GA signaling by inhibiting RGA function. CDK8, but not the kinase-dead mutant (CDK8^D176A^)^56^, rescued the *cdk8* hypocotyl response to GA (**Figures 4A, 4B**, **S3A and S3B**), indicating that CDK8 kinase activity is essential for CDK8-induced GA response.

**Figure 4.**
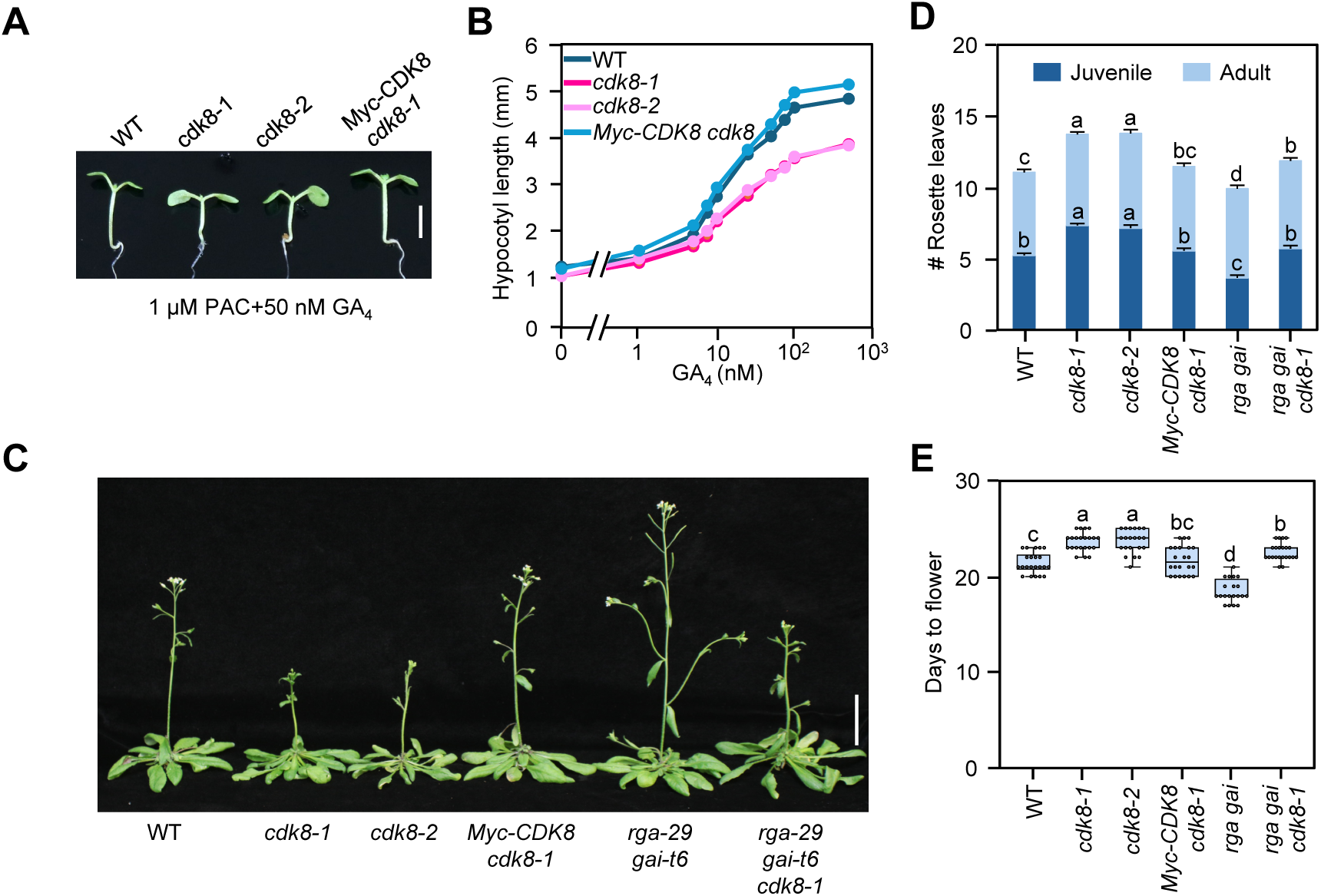
CDK8 reduced RGA function to promote GA signaling. (A)-(B) the *cdk8* mutants displayed a reduced GA response in hypocotyl growth. Seedlings were grown in medium containing 1 μM PAC and varying concentrations of GA_4_. The photo was taken and hypocotyl lengths were measured at day 9. In (A), Bar = 3 mm. In (B), Average hypocotyl lengths. Means ± SE. n =11. (C)-(E), the *cdk8* mutants showed delayed phase transitions in leaf development and flowering time, and *rga-29 gai-t6* partially rescued these phenotypes. In (C), photo showing representative 31d-old plants grown under LD. Bar = 3 cm. (D) Average rosette leaf numbers after bolting. Juvenile leaves are rosette leaves that lack abaxial trichomes. The values plotted are the means ± SE. n =10-11. (E) Days to flower of different lines are shown in the boxplot. n=10-11. Center lines and box edges are medians and the lower/upper quartiles, respectively. Whiskers extend to the lowest and highest data points within 1.5× interquartile range (IQR) below and above the lower and upper quartiles, respectively. In (D)-(E), different letters above the bars represent significant differences [*p* < 0.01 for juvenile leaves and *p* < 0.05 for rosette leaf numbers in (D), *p* < 0.05 in (E)] as determined by two-tailed Tukey’s HSD test. The phenotypic analysis was repeated two times with similar results.

At the adult stage, both *cdk8* mutants exhibited delayed vegetative and reproductive phase transitions compared with WT (**Figures 4C-4E**). To examine the genetic interaction between *CDK8* and two *DELLAs* (*RGA* and *GAI*), the *cdk8-1 rga-29 gai-t6* triple mutant was generated, in which the *rga-29* and *gai-t6* null alleles are caused by T-DNA and transposon Ds insertions in their respective coding regions^7,57^. We found that *rga-29 gai-t*6 partially rescued hypocotyl growth, juvenile-to-adult leaf transition and flowering time of *cdk8-1* (**Figures 4C-4E**), consistent with the notion that CDK8 promotes GA signaling by reducing RGA and GAI activity.

### CDK8-mediated phosphorylation of S170 reduced RGA activity

To identify CDK8-mediated phosphosite(s) in RGA, we performed liquid chromatography-electrospray ionization-tandem mass spectrometry (LC-ESI-MS/MS) using affinity-purified His-FLAG-RGA^GKG^ protein from transgenic *Arabidopsis* Col-0 ecotype plants carrying *P_RGA_:His-FLAG-RGA^GKG^* in *ga1-13* and *ga1-13 cdk8-1* backgrounds, separately. Relative peptide abundance was estimated by semi-quantitative analysis of ion currents detected in MS1 survey scans. All RGA phosphorylation sites detected were located within the PolyS and PolyS/T regions flanking the DELLA domain: LSN-peptide [Pep1, LSNHGTSSSSSSISK(DK), 5% phosphorylation] and LKS-peptide [Pep2, (LK)SCSSPDSMVTSTSTGTQIGK, 32.9%] (**Figures 5A, 5B and S4, Table S3**). These phosphosites largely match those identified in our previous MS analysis of FLAG-RGA^GKG^ purified from *ga1-3* in the L*er* ecotype background^42^. However, Pep1 phosphorylation was higher in L*er* than in Col-0 ecotype (30.5% vs. 5.0%). Additionally, only S170 phosphorylation within Pep2 was detected in Col-0, whereas multiple phosphosites were identified in the L*er* background.

**Figure 5.**
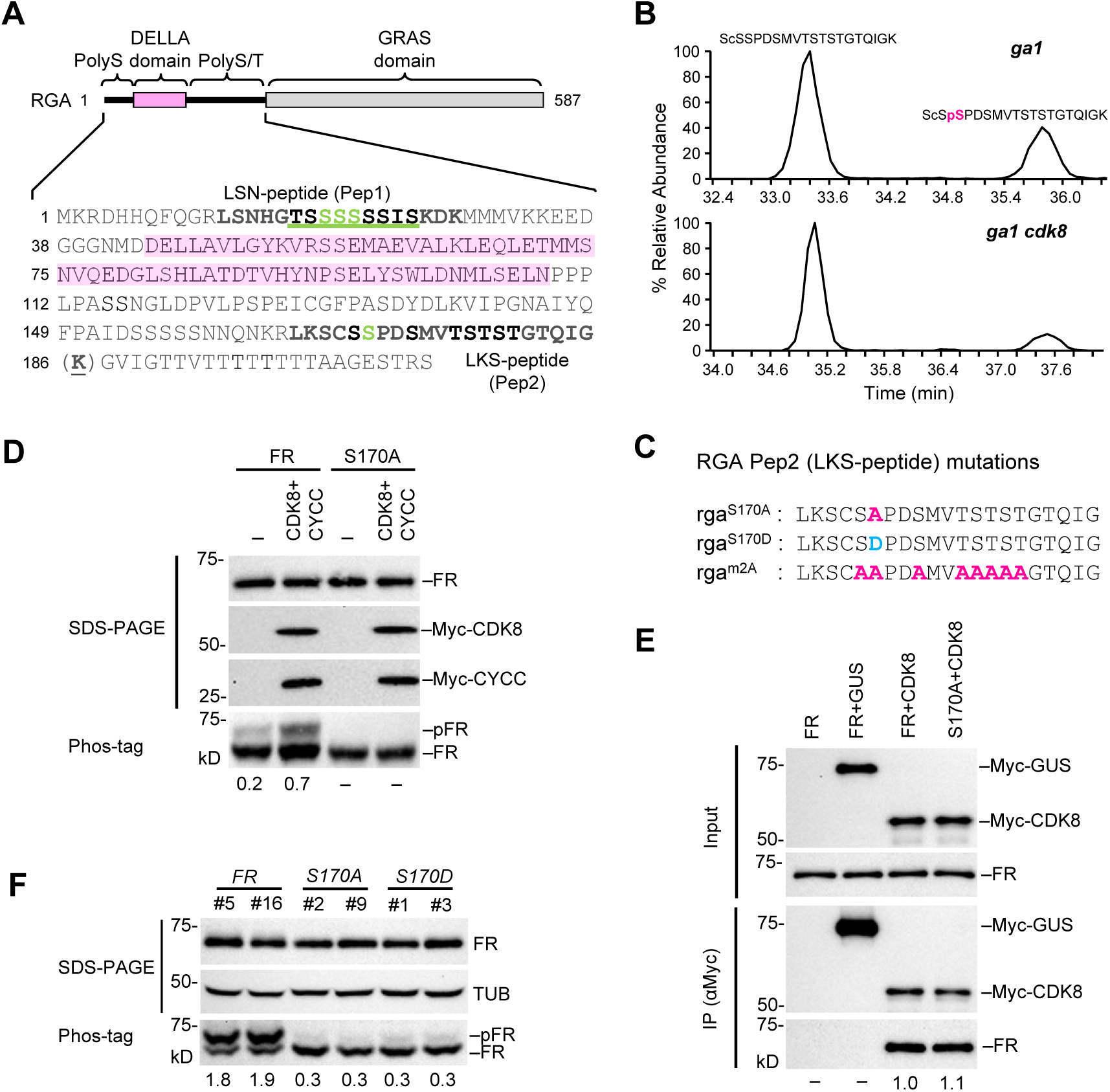
Identification of CDK8-mediated phosphosite in RGA. (A) RGA phosphorylation sites identified by MS/MS analysis. The schematic shows the RGA protein structure. Two disordered regions are depicted as solid black lines. The DELLA domain is shaded in pink. The LSN-peptide (Pep1) and LKS-peptide (Pep2) containing phosphorylation are in black bold letters. The S/T residues in green letters are confirmed phosphorylation sites. The underlined residues in Pep1 are amino acid stretches, in which one or more residues are modified (in addition to the identified sites), but the specific residues could not be mapped. The underlined K in parenthesis indicates the extra Lys residue in RGA^GKG^ for creating an additional trypsin cleavage site. (B) MS1 extracted ion chromatogram (XIC) showing reduced S170 phosphorylation within RGA Pep2 in *cdk8*. Relative XICs of pS170 from *Arabidopsis ga1-13* (top) and *ga1-13 cdk8-1* (bottom). Red “pS” denotes the phosphorylated serine; lowercase “c” denotes alkylated cysteine. (C) Mutated residues in rga^S170A^, rga^S170D^ and rga^m2A^ are highlighted in red (S/T-to-A) or blue (S-to-D). (D) S170A abolished CDK8/CYCC-mediated RGA phosphorylation in *N. benthamiana*. (E) Co-IP assays showing that S170A did not affect Myc-CDK8 binding in *N. benthamiana*. The amount of FLAG-RGA pulled down by Myc-CDK8 was set as 1.0. –, not detectable. Myc-GUS was included as a negative control. (F) Expression and phosphorylation pattern of *P_RGA_:His-FLAG-RGA/-rga^S170A^/rga^S170D^ ga1 dP* transgenic lines using standard SDS-PAGE and Phos-tag gels. Protein blots were probed with anti-FLAG antibody or anti-tubulin (TUB, as a loading control). Representative images of 2 biological repeats are shown. In (D) and (F), the ratios of pFR/unphosphorylated FR are shown below the Phos-tag gel blot.

Importantly, the *cdk8-1* mutation reduced relative phosphorylation abundance of S170 within Pep2 by 2-fold (32.9% in *ga1* vs 16.0% in *ga1 cdk8*) (**Figure 5B**), while phosphorylation within Pep1 was unchanged (**Table S3**). To test whether S170 in RGA is the primary CDK8-dependent phosphosite, we expressed wild-type RGA or the mutant rga^S170A^, either alone or together with CDK8 and CYCC, in *N. benthamiana*. S170A is expected to block phosphorylation at S170. Phos-tag gel analysis showed that the S170A mutation abolished CDK8-mediated RGA phosphorylation (**Figure 5C and 5D**), although co-IP assays indicated that this mutation did not affect CDK8-RGA interaction (**Figure 5E**). These results indicate that S170 is the major CDK8 phosphosite in RGA.

To investigate how S170 phosphorylation affects RGA function in planta, we generated transgenic *Arabidopsis* in the *ga1 dP* background carrying either *P_RGA_:His-FLAG-rga^S170A^* or *- rga^S170D^*. While the S170A substitution abolishes phosphorylation at this site, S170D serves as a phosphomimetic substitution. Three independent homozygous lines for each construct, with expression levels comparable to the previously reported *P_RGA_:His-FLAG-RGA* lines^24^, were selected for phenotypic analysis (**Figure S5A**). Phos-tag gel analysis showed that the S170A mutation reduced overall RGA phosphorylation to ∼16% of the wild-type level (**Figure 5F**). As expected, *His-FLAG-RGA* restored the dwarf phenotype in *ga1 dP* plants. Notably, the phosphomimetic S170D substitution slightly decreased RGA activity in repressing stem growth and floral induction (**Figures 6A, 6B and S5B-SD**), supporting the idea that CDK8-mediated phosphorylation of S170 attenuates RGA activity. In contrast, S170A did not further reduce growth compared with wild-type RGA, possibly because the control RGA lines already display an extremely dwarf phenotype. Consistent with these observations, hypocotyl elongation assays showed that S170D enhanced the GA response in hypocotyl growth (**Figures 6C and 6D**). Together, these results indicate that CDK8 promotes GA signaling by down-regulating RGA activity through phosphorylation of S170.

**Figure 6.**
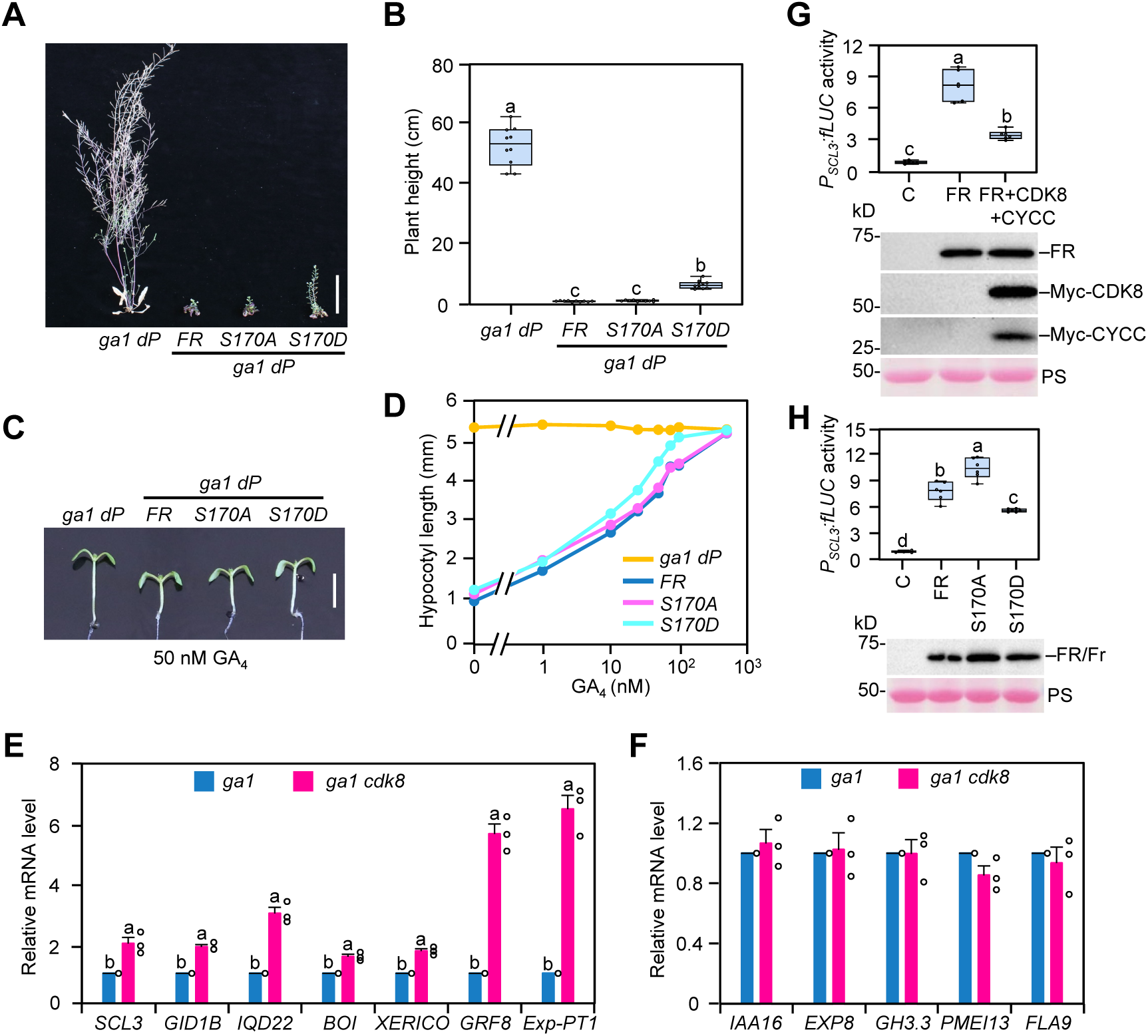
Phosphomimetic S170D reduced RGA function in planta. (A)-(B) Phenotypes of *P_RGA_:His-FLAG-RGA/rga* transgenic lines in the *ga1 dP* background. *FR*, line #16; *S170A*, line #9); S170D, line #1. In (A), Representative 93d-old plants grown under LD. Bar = 5 cm. In (B), Boxplot showing heights of different lines as labeled. n=10. (C)-(D) *His-FLAG-rga^S170D^ ga1 dP* displayed an enhanced GA response in hypocotyl growth. Seedlings were grown in medium containing varying concentrations of GA_4_. Hypocotyl lengths were measured at day 9. Bar = 3 mm. In (D), Average hypocotyl lengths are means ± SE. n =12. (E)-(F) RT-qPCR analyses showing *cdk8* caused elevated expression of RGA-induced genes (E), but did not change expression of RGA-repressed genes significantly (F, *p* > 0.05). *PP2A* was used to normalize different samples. Means ± SE of three biological replicates are shown. In e, Different letters above the bars represent significant differences (*p* < 0.01) by two-tailed Student’s *t*-test. (G)-(H) Dual LUC assays in *N. benthamiana*. Top panel: the boxplot shows relative fLUC activity. C, empty effector as a negative control. Lower panel: immunoblots shows expression levels of effector proteins in different samples as detected by anti-FLAG and anti-Myc antibodies. PS, Ponceau staining for even loading. In (G), co-expression of CDK8/CYCC reduced RGA-induced *SCL3* transcription. In (H), RGA activity was elevated by S170A, but reduced by S170D substitution. In all boxplots shown in (B) and (G-H), center lines and box edges are medians and the lower/upper quartiles, respectively. Whiskers extend to the lowest and highest data points within 1.5× interquartile range (IQR) below and above the lower and upper quartiles, respectively. Different letters above the bars represent significant differences (*p* < 0.01) as determined by two-tailed Tukey’s HSD test. All assays in (A-D, G-H) were repeated two times with similar results.

Interestingly, this inhibitory effect of CDK8 on RGA activity contrasts with our prior finding that substituting eight S/T-to-A residues in RGA Pep2 (the m2A mutant) reduced RGA function (**Figure 5C**), suggesting that phosphorylation within Pep2 can enhance RGA activity^42^. We therefore propose that CDK8 and another, as yet unidentified, protein kinase may regulate RGA differently by targeting distinct phosphorylation sites. While CDK8-mediated phosphorylation of S170 reduces RGA activity, phosphorylation of other S/T residues within Pep2 may enhance its function.

### CDK8 does not affect RGA stability or its interaction with PIF3, BZR1 or H2A

To understand how CDK8-mediated phosphorylation reduces RGA function, we examined whether *cdk8* affects RGA subcellular localization or stability in response to GA. The *cdk8* mutation did not alter GA-induced RGA degradation (**Figure S6A**) or its nuclear localization (**Figure S6B**). RT-qPCR analysis showed that *cdk8* mutation increased expression of RGA-induced genes (**Figure 6E**) but did not affect RGA-repressed genes (**Figure 6F**), suggesting that CDK8 mainly suppresses the transactivation activity of RGA. To further test this idea, we performed dual luciferase (LUC) assays using the transient expression system in *N. benthamiana*. The *P_SCL3_:firefly LUC (fLUC)* was used as the reporter because *SCL3* is a direct DELLA target whose transcription is activated by DELLA^58,59^. *35S:Renilla LUC (rLUC)* served as an internal control to normalize transformation efficiency. Co-expression of CDK8 and CYCC reduced RGA-induced *P_SCL3_:fLUC* expression (**Figure 6G)**, indicating that CDK8 phosphorylation suppresses RGA activity. Consistently, the S170A mutation enhanced RGA activity, whereas the phosphomimetic S170D reduced it (**Figure 6H**), supporting that CDK8-mediated phosphorylation of S170 attenuates RGA function.

Because formation of TF-RGA-H2A complexes at target chromatin is essential for RGA function^24^, we next examined whether CDK8 affects RGA interactions with TFs or H2A. In vitro pulldown assays were performed using recombinant GST-PIF3 and GST-BZR1 together with protein extracts from transgenic *Arabidopsis* expressing FLAG-RGA^GKG^ in either the WT or *cdk8-1* background. GST-PIF3 and GST-BZR1 pulled down similar amounts of His-FLAG-RGA from both backgrounds (**Figures 7A and S7**). Co-IP assay using FLAG-RGA^GKG^ transgenic lines also showed that the *cdk8* mutation did not affect the RGA–H2A interaction in planta (**Figure 7B**). Similarly, the S170A and S170D substitutions did not alter RGA binding to H2A. In contrast, the previously described m2A mutation (eight S/T-to-A substitutions in Pep2) markedly weakened the RGA–H2A interaction (**Figure 7C**), consistent with our earlier findings^42^.

**Figure 7.**
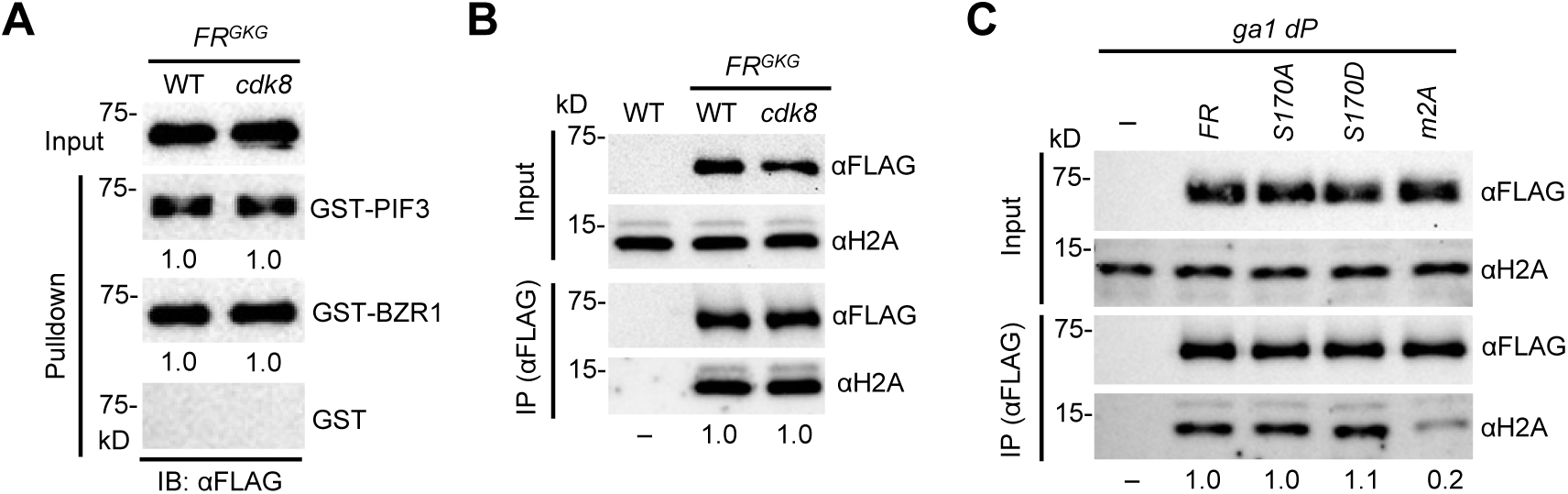
*cdk8* did not alter RGA binding to PIF3, BZR1 or H2A. (A) In vitro pulldown assays showing that *cdk8-1* did not affect RGA binding to GST-PIF3 or GST-BZR1. Recombinant GST, GST-PIF3 and GST-BZR1 bound to glutathione–Sepharose beads were used separately to pull down His-FLAG-RGA^GKG^ (FR^GKG^) from protein extracts of 10d-old *P_RGA_-His-FLAG-RGA^GKG^* transgenic *Arabidopsis* in WT or *cdk8-1* background that were grown in media containing 1 μM PAC. Immunoblots containing input *Arabidopsis* extracts and pulldown samples were detected with an anti-FLAG antibody. Ponceau S-stained blots indicated that similar amounts of the GST/GST-fusion proteins were used in all pulldown assays (**Figure S7**). Relative amounts of FR^GKG^ pulled down by GST fusion proteins are shown below each blot with the FR^GKG^ levels in the WT pulldown sample set as 1.0. (B)-(C) Co-IP assays showing that *cdk8-1* did not alter RGA-H2A binding (B), and that *m2A* significantly reduced endogenous H2A binding, but *S170A* or *S170D* did not (C). In (B), FR^GKG^ from protein extracts of transgenic lines prepared as described in (A) was immunoprecipitated using anti-FLAG agarose. WT Col-0 was included as a negative control. In (C), His-FLAG-RGA/rga from protein extracts of transgenic *Arabidopsis* in *ga1 dP* background carrying *P_RGA_:His-FLAG-RGA/rga* were immunoprecipitated using anti-FLAG agarose. Immunoblots containing input *Arabidopsis* extracts and immunoprecipitated samples were detected with anti-FLAG and anti-H2A antibodies, separately. Representative images of two biological repeats are shown. Relative amounts of H2A co-immunoprecipitated with FLAG-RGA/rga are shown below the blots. –, not detectable. In (A-C), representative images of two biological repeats are shown.

### CDK8 attenuates RGA transactivation by weakening RGA-MED15 interaction

The coactivator MED15 interacts with RGA to promote transcription of RGA-induced genes^25,60^. Because MED15 and several other Mediator subunits were identified in the RGA proximitome (**Figure 1C**), we tested whether CDK8/CYCC affects RGA–MED15 interaction. Yeast three-hybrid (Y3H) assays were performed using Gal4 DNA-binding domain (DB)-MED15, Gal4 transactivation domain (AD)-RGA in the absence or presence of Myc-CDK8/Myc-CYCC (**Figures 8A** and **S8**). Myc-CDK8^D176A^ served as a kinase-dead control. Interaction strength was assessed by growth on His^−^ media containing increasing concentrations of the competitive HIS3 inhibitor 3-aminotriazole (3-AT). Co-expression of CDK8 and CYCC weakened MED15-RGA interaction, as indicated by yeast growth only in the absence of 3-AT rather than at 10 mM. In contrast, CDK8^D176A^ had no effect, indicating that CDK8 kinase activity is required.

**Figure 8.**
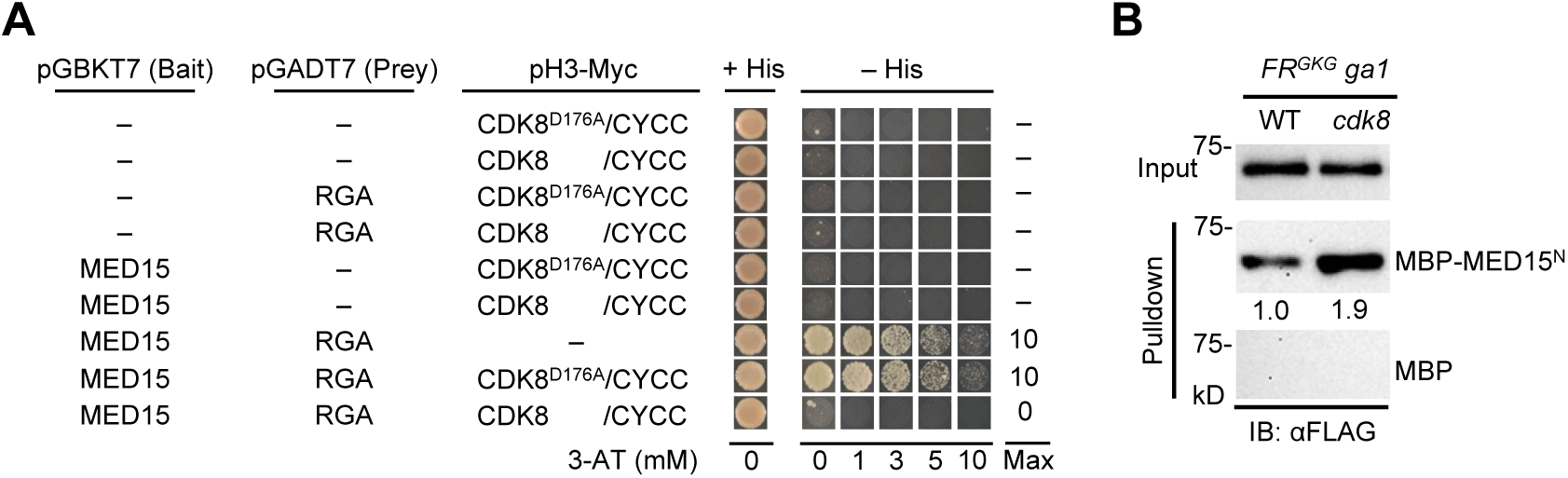
CKM weakened RGA-MED15 interaction by Y3H and pulldown assays. (A) Y3H assay showing CDK8/CYCC reduced RGA-MED15 interaction. Full-length MED15 was fused to the Gal4 DB domain as the bait, and the full-length RGA was fused to Gal4 AD domain as the prey. CDK8, CDK8^D176A^ and CYCC were expressed using pH3-Myc vector. Immunoblot analyses showed similar levels of RGA and CDK8 proteins across samples (**Figure S8**). Interaction of DB and AD fusion proteins in the PJ69-4A yeast cells was scored by the relative growth in –His media containing varying concentrations of 3-AT as labeled. – indicates empty vector, or no growth at 0 mM 3-AT. Max, detectable cell growth at maximum 3-AT concentration. (B) In vitro pulldown assay showing FLAG-RGA^GKG^ from the *cdk8-1* background binds to MBP-MED15^N^ more strongly than that from WT *CDK8* background. Recombinant MBP, and MBP-MED15^N^ bound to amylose beads were used separately to pull down FR^GKG^ from protein extracts of 10d-old *P_RGA_:His-FLAG-RGA^GKG^* transgenic *Arabidopsis* in *ga1* or *ga1 cdk8-1* background. Immunoblots containing input *Arabidopsis* extracts and pulldown samples were detected with an anti-FLAG antibody. Ponceau S-stained blots indicated that similar amounts of the MBP/MBP-fusion proteins were used in the pulldown assays (**Figure S11B**). In (A-B), two biological repeats showed similar results.

A limitation of these Y3H assays was that full-length (FL) MED15 protein was not detectable by immunoblotting and was also unstable when expressed in *N. benthamiana*. To address this, we mapped RGA-interacting region of MED15 using co-IP and Y2H assays. Three MED15 fragments were detectable and capable of binding RGA, but the N-terminal region (amino acids 1–450, MED15^N^), which contains the KIX domain, showed the strongest interaction (**Figures S9 and S10**). The KIX domain is necessary and sufficient for RGA binding, although weaker than MED15^N^ (**Figure S10**).

The inhibitory effect of CDK8 on MED15-RGA interaction was further verified by co-IP and in vitro pulldown assays using MED15^N^. For the Co-IP assay, FLAG-RGA was transiently expressed alone or together with YFP-MED15^N^ in *N. benthamiana* leaves, with or without Myc-CDK8 and Myc-CYCC. YFP-GUS and Myc-CDK8^D176A^ were included as negative controls. FLAG-RGA co-immunoprecipitated more strongly with YFP-MED15^N^ in the absence of CDK8/CYCC than when CDK8/CYCC were co-expressed (**Figures 9 and S11A**). Consistently, CDK8^D176A^ did not weaken RGA-MED15^N^ interaction. In vitro pulldown assays using recombinant MBP-MED15^N^ and protein extracts from *Arabidopsis* carrying *P_RGA_:His-FLAG-RGA^GKG^* in either *ga1-13* or *ga1-13 cdk8-1* backgrounds further showed that the *cdk8-1* mutation increased MED15^N^–RGA binding (**Figures 8B and S11B**).

**Figure 9.**
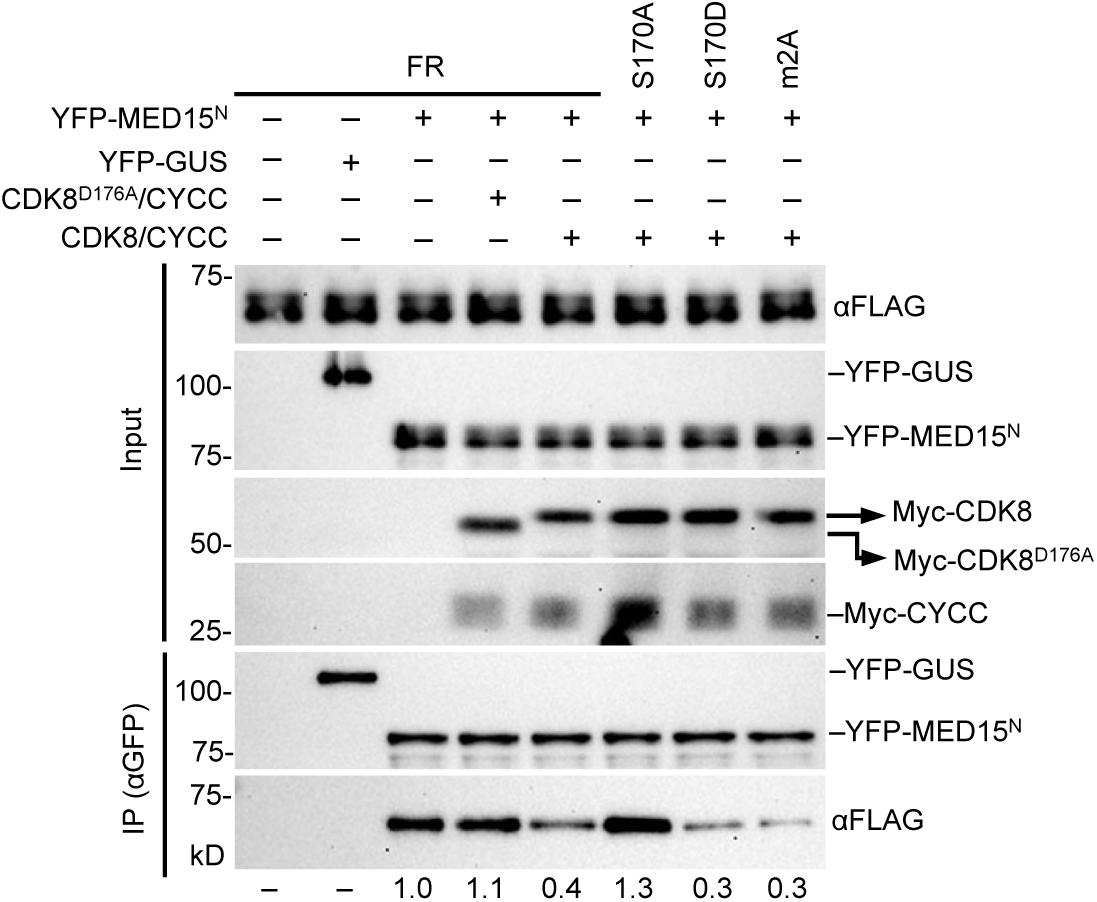
Co-IP assays showed that CDK8/CYCC reduced RGA-MED15^N^ interaction, and that S170D and S170A substitutions reduced and enhanced MED15^N^ binding, respectively, in *N. benthamiana.* FLAG-RGA/rga was expressed alone or together with YFP-MED15^N^ in the presence or absence of Myc-CDK8/-CYCC. YFP-GUS and Myc-CDK8^D176A^ were included as negative controls. Levels of phosphorylated FLAG-RGA/rga in input samples were examined by Phos-tag gel analysis (**Figure S11A**). YFP-MED15^N^ or -GUS were immunoprecipitated using anti-GFP agarose. Immunoblots containing total protein extracts (input) or immunoprecipitated samples were probed with αFLAG, αMyc or αGFP. Relative amounts of FLAG-RGA/rga co-immunoprecipitated with YFP-MED15^N^ are shown below the blot. –, not detectable. Representative images of 3 biological repeats are shown.

To examine directly the role of S170 phosphorylation in RGA-MED15 interaction, co-IP assays were performed in *N. benthamiana* expressing FLAG-RGA or rga^S170A^ or rga^S170D^ together with YFP-MED15^N^, with or without CDK8/CYCC. The phosphomimetic S170D mutation reduced MED15 binding, whereas S170A increased binding compared with wild-type RGA (**Figures 9 and S11A**). We also examined the m2A mutant (eight S/T-to-A substitutions in Pep2). In contrast to S170A, m2A weakened MED15 binding. These results suggest that phosphorylation of S170 decreases RGA activity by weakening MED15 interaction, whereas phosphorylation at other S/T residues within Pep2 enhances RGA function.

## DISCUSSION

Based on our findings, we propose a working model in which the CDK8 kinase module (CKM) functions as a regulatory switch that modulates DELLA-dependent transcription and thereby fine-tunes GA responses (**Figure 10**). In this model, CKM attenuates the transactivation capacity of the DELLA protein RGA by limiting recruitment of the coactivator Mediator complex to GA-responsive promoters. Mediator is a conserved transcriptional hub that transmits regulatory inputs from transcription factors (TFs) to RNA polymerase II (Pol II). The core Mediator (cMED) complex contains head, middle, and tail modules that interact with Pol II and TFs, respectively. Association of the CDK8 kinase module—composed of CDK8, Cyclin C (CYCC), MED12, and MED13—adds a dynamic regulatory layer to Mediator function^48,49^. Although initially characterized as a transcriptional repressor that blocks Mediator–Pol II interaction, CKM is now recognized as a context-dependent regulator capable of either repressing or activating transcription through CDK8 kinase activity in response to developmental and stress signals^48,61^. In *Arabidopsis*, CDK8 has been implicated in ABA signaling during drought stress^62^ and in salicylic acid–mediated defense responses^63,64^, but its connection to GA/DELLA signaling had not been established.

**Figure 10.**
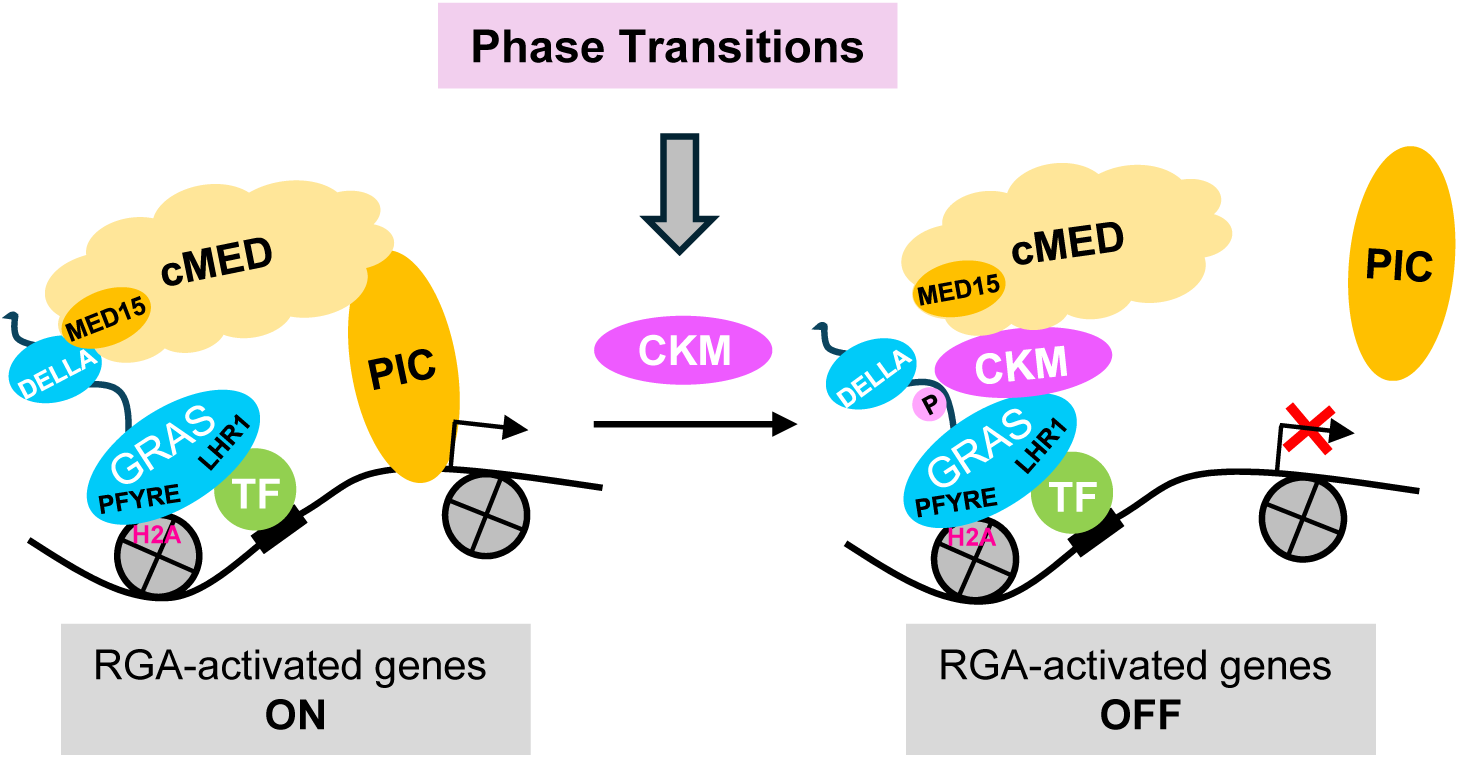
Working model of CKM-mediated attenuation of DELLA function. DELLA proteins are recruited to target chromatin through interactions with TFs, forming TF–DELLA–H2A complexes that regulate the transcription of GA-responsive genes. CDK8 kinase module (CKM) promotes GA-induced vegetative and reproductive phase transitions by phosphorylation of RGA at Ser170 within the disordered PolyS/T region between the DELLA and GRAS domains. This modification weakens RGA-MED15 interaction, thereby limiting core Mediator complex (cMED) recruitment to RGA-activated and GA-repressed promoters. PIC, RNA polymerase II preinitiation complex.

Previous studies showed that DELLAs are recruited to target chromatin through interactions with transcription factors (TFs), forming TF–DELLA–H2A complexes that regulate the transcription of GA-responsive genes^42^. DELLA-mediated transcriptional activation further depends on recruitment of the Mediator coactivator complex via interaction with the tail subunit MED15 ^25^. Building on this framework, we propose that CKM associates with RGA-containing transcriptional complexes during vegetative and reproductive phase transitions, where CDK8 phosphorylates RGA at Ser170. This modification weakens the interaction between RGA and MED15, thereby reducing Mediator recruitment and dampening transcriptional activation of RGA-induced genes.

This model is supported by multiple lines of evidence. TurboID proximity labeling identified CDK8 together with MED12 and MED13 in the RGA proximity proteome. Phos-tag gel and MS/MS analyses demonstrated CKM-mediated phosphorylation of RGA at Ser170 in planta. Functional assays further showed that CKM weakens RGA–MED15 association in both Y3H systems and co-IP assays. Consistent with this mechanism, *cdk8* and *rga gai* mutants exhibit opposing phenotypes during vegetative and reproductive phase transitions—developmental processes that are promoted by GA signaling and repressed by DELLAs^65,66^. In addition, *MED12* and *MED13* have been reported to promote these developmental transitions^67–69^. Together, these findings support a role for CKM in promoting GA signaling by limiting DELLA-mediated transcription through phosphorylation-dependent control of Mediator recruitment.

Our results further indicate that RGA activity is modulated by multiple phosphorylation sites within the Pep2 region. Ser170 lies within the intrinsically disordered PolyS/T region between the DELLA and GRAS domains. At first glance, the inhibitory effect of Ser170 phosphorylation appears inconsistent with our previous observation that substitution of eight Ser/Thr residues within Pep2, including S170 (the m2A mutant, **Figure 5C**), reduces RGA activity in planta^42^, suggesting that phosphorylation within this region promotes RGA function. However, the underlying phenotypes reflect distinct molecular effects. The S170A mutation enhances RGA–MED15 interaction without affecting RGA binding to histone H2A, whereas the m2A mutation strongly disrupts the RGA–H2A interaction^42^ (**Figure 7C**). These findings suggest that phosphorylation at different Pep2 residues selectively modulates RGA interactions with distinct cofactors. We therefore propose that RGA activity is dynamically tuned by multisite phosphorylation within Pep2, potentially mediated by different kinases with opposing effects. In this model, CDK8 phosphorylation of Ser170 attenuates RGA activity by limiting Mediator recruitment, whereas phosphorylation of other Pep2 residues enhances RGA function, at least in part by stabilizing interaction with H2A.

The functional significance of DELLA phosphorylation has long been debated, with studies proposing roles in DELLA degradation, stabilization, or activity. Our results demonstrate that phosphorylation can regulate DELLA transcriptional activity independently of protein stability and that distinct phosphorylation events within intrinsically disordered regions (IDRs) can produce different regulatory outcomes. These findings are consistent with the emerging concept that IDRs serve as regulatory hubs enriched in clustered phosphorylation sites, enabling context-dependent control by tuning the specificity and affinity of protein-protein interactions rather than functioning as simple on-off switches ^70–72^.

Beyond the CDK8 kinase module highlighted here, our TbID proximity labeling analysis also uncovered a large set of previously unrecognized RGA-proximal proteins linked to transcriptional regulation. These findings markedly expand the known RGA interaction landscape and suggest that RGA engages a broad transcriptional regulatory network comprising transcription factors, transcriptional cofactors, chromatin remodelers, and histone acetyltransferase complexes. This expanded RGA proximitome reveals strong connections between RGA and core transcriptional regulatory machinery, providing a framework for mechanistic dissection of RGA-mediated transcriptional control.

Taken together, our work uncovers a mechanism linking the Mediator kinase module to GA signaling through phosphorylation-dependent control of DELLA activity. By modulating the interaction between RGA and Mediator, CKM provides a tunable layer of transcription regulation that connects kinase signaling with hormone-responsive gene expression. This mechanism highlights how phosphorylation within IDRs can dynamically shape transcriptional regulatory networks during plant development.

## Limitations of the study

While our study shows that distinct phosphorylation events within intrinsically disordered regions (IDRs) of the DELLA protein RGA can lead to different regulatory outcomes, several mechanistic questions remain unresolved. How phosphorylation of Ser170 modulates RGA–MED15 binding, and how phosphorylation of other serine residues within Pep2 influences the RGA–H2A interaction, are not yet understood. In addition, the kinase(s) responsible for phosphorylating other Pep2 residues remain unidentified, and how these phosphorylation events are integrated to regulate DELLA activity during plant development remains unclear.

## Methods

### Plant materials, growth conditions, and generation of transgenic lines

In most experiments, *Arabidopsis* plants were grown in the growth room under long-day (LD) conditions with 145 µmol m^-2^ s^-1^ light intensity (16 h light, 8 h dark, 22 °C). For dim light treatment, seedlings were grown in 16 µmol m^-2^ s^-1^ light intensity under LD for hypocotyl elongation assays or in 10 µmol m^-2^ s^-1^ light intensity under SD (8 h light) at 22 °C for RT-qPCR analysis in Fig. 5e-f. All Arabidopsis lines used in this study are in the Col-0 ecotype background. The *ga1-13* (SALK_109115), *ga1-13 dQ* (quadruple *della* with *RGA*), *ga1-13 della pentuple* [(*ga1 dP; dP contains rga-29*: SALK_089146*; gai-t6* (backcrossed 6x with Col-0), *rgl1:* SALK_136162*; rgl2:* SALK_027654*; rgl3-3:* CS16355)], and *rga-29 gai-t6* were described previously^24,57,73^. The *cdk8-1* (SALK_138675)^56^ and *cdk8-2* (SALK_016169)^56^ were obtained from Dr. Clint Chapple at Purdue University. The *med15-4* (SAIL_792_F02, a.k.a. *nrb4-4*)^74^ mutant was obtained from the Arabidopsis Biological Resource Center (https://abrc.osu.edu/). The triple homozygous mutant *rga-29 gai-t6 cdk8-1* were generated by crossing *rga-29 gai-t6* with *cdk8-1*. The double homozygous mutant *ga1-13 cdk8-1* was generated by crossing *ga1-13* with *cdk8-1*. Transgenic lines *P_RGA_:His-FLAG-RGA ga1 dP* and *P_RGA_:His-FLAG-RGA^m2A^ ga1 dP* were reported previously^24,42^. Transgenic Arabidopsis lines*, P_RGA_:His-FLAG-RGA^GKG^* Col-0, *P_RGA_:His-FLAG-RGA-TurboID ga1 dP*, *P_RGA_:His-FLAG-NLS-GFP-TurboID ga1 dQ, P_RGA_:His-FLAG-RGA^S170A^ ga1 dP,* and *P_RGA_:His-FLAG-RGA^S170D^ ga1 dP,* were generated by introducing each construct into specific genetic background by agrobacterium-mediated transformation. Independent transgenic lines with single insertion were selected by Basta resistance. Multiple independent transgenic lines (4 to 9) for each construct were screened by immunoblot analysis to select for lines that expressed His-FLAG-RGA/-rga/RGA-TurboID or GFP-TurboID protein at similar levels. *P_RGA_:His-FLAG-RGA^GKG^ ga1-13, P_RGA_:His-FLAG-RGA^GKG^ cdk8-1,* and *P_RGA_:His-FLAG-RGA^GKG^ ga1-13 cdk8-1* were generated by crosses between *P_RGA_:His-FLAG-RGA^GKG^* Col-0 and *ga1-13 cdk8-1*. *35S:Myc-CDK8* and *35S:Myc-CDK8^D176A^* were transformed into *cdk8-1* mutant separately to generate Basta-resistant transgenic lines (3 and 8 lines each)*. P_RGA_:His-FLAG-RGA ga1 dP* line #5 was transformed with *P_Ubi10_:Myc-CDK8* to generate 3 double transgenic lines. Primers for genotyping are listed in **Table S4**.

### Plasmid construction

The following plasmids were described previously: *P_SCL3_:fLUC*, and *35S:rLUC*^75^ for dual LUC assays, pEG3F-RGA/rga-m2A/NT3/CT3 (35S:FLAG-RGA/m2A/NT3/CT3)^42^, pCR8GW-GUS-NLS^59^, and pEG203-H2A (35S:Myc-H2A)^24^ for transient expression in *N. benthamiana*, pBm43GW^76^ for cloning, GST-PIF3, GST-BZR1^55^, and GST-H2A^24^ for expression in *E. coli,* and pH3-Myc-GW^77^ for Y3H assays. Primers and plasmid constructs are listed in **Tables S4 and S5**, respectively. All new DNA constructs generated from PCR amplification were sequenced to ensure that no mutations were introduced. Construction of pRGA-His-3xFLAG-RGA-TurboID/NLS-GFP-TurboID/RGA^S170A^/RGA^S170D^ were generated using four constructs: J035 (RGA promoter, 8.1kb), pBm43GW^76^, J015(RGA 3’UTR) and pDONR207-His-3xFLAG-RGA-TurboID/NLS-GFP-TurboID/RGA^S170A^/RGA^S170D^ by Gateway LR reaction.

### GA response assays and adult phenotype analyses

For hypocotyl elongation analysis of WT Col-0 and *cdk8* mutants, surface-sterilized seeds were imbibed in water for 3 days at 4 °C, then plated on 0.5x Murashige and Skoog (MS) medium with 0.5% sucrose, supplemented with 1 μM paclobutrazol (PAC, a GA biosynthesis inhibitor), and varying concentrations of GA_4_ for 9 days under LD conditions with dim light (16 µmol m^-2^ s^-1^ white light). For *ga1 dP* and transgenic lines carrying *P_RGA_:His-FLAG-RGA/RGA^S170A^/RGA^S170D^* in the *ga1 dP* background, the seeds were imbibed in 10 μM GA_4_ to promote germination and then washed thoroughly with water before plating on 0.5x MS medium with 0.5% sucrose and varying concentrations of GA_4._ Hypocotyl lengths were measured using ImageJ software (http://rsb.info.nih.gov/ij).

To characterize the adult phenotypes, the seeds were imbibed similarly as described above and then were sown in soil under LD. Phenotype measurements were performed as described^73^. Days to flower were measured as number of days from sowing until floral buds are clearly visible. The experiment was repeated 2 times for all with similar results.

### Statistical analyses

Statistical analyses for all quantitative data were conducted using Student’s t-test for comparisons between two samples, or one-way ANOVA followed by Tukey’s Honestly Significant Difference (HSD) mean separation test for comparisons among multiple samples, using SPSS Statistics 17.0 software.

### Yeast two-hybrid and yeast three-hybrid Assays

The yeast strain PJ69-4A and the Matchmaker Gold Yeast Two-Hybrid System (Clontech, Palo Alto, CA) were used, in which the bait vector pGBKT7 contains the Gal4 DNA-binding domain and the prey vector pGADT7 with Gal4 activation domain. For Y2H assays, yeast transformation and 3-amino-1,2,4-triazole (3-AT) tests were performed as described previously^55^. Varying concentrations of 3-AT were included in medium lacking Leu, Trp, and His. For each combination, 3 μL yeast cells with OD_600_ values of 0.4, respectively, were spotted on media plates. For Y3H assays, a third vector pH3-Myc-GW was used to express the third set of proteins, and the protein-protein interaction analysis on media minus Leu, Trp, Ura and His with different 3-AT concentrations.

### Transient expression and dual luciferase assay in *Nicotiana benthamiana*

Transient expression of various epitope-tagged proteins in *N. benthamiana* was performed as described^75^. The *N. benthamiana* leaves were harvested 48 h after Agro-infiltration for protein extraction, followed by dual luciferase assays using dual-luciferase reporter assay system (Promega). Three biological repeats were conducted for each effector combination.

### Reverse transcription (RT)-quantitative PCR (qPCR) and immunoblot analyses

Total Arabidopsis RNA extraction and RT-qPCR analysis were performed as described^42^. Relative transcript levels were determined by normalizing with *PP2A* (AT1G13320)^78^. Primers for the qPCR are listed in **Table S4**.

For immunoblot assays, total proteins were extracted from 10 days–old seedlings using the 4% SDS extraction buffer as described previously. Immunoblot analyses were performed using horseradish peroxidase (HRP)-conjugated anti-FLAG M2 mouse monoclonal (Sigma Aldrich A8592, 1:10,000 dilution), rat anti-RGA antiserum (DU18, 1:1,000)^15^, rabbit anti-H2A monoclonal antibody (Abcam ab177308, 1:1,000 dilution), rabbit anti-H3 polyclonal antibody (Abcam ab1791, 1:5,000), mouse anti-tubulin antibody (Sigma T5168, 1:200,000 for **Figures 5F and S5A;** 1:100,000 for **Figure S6B**), mouse anti-GFP antibody (Roche 11814460001, 1:2,000 dilution for input samples or 1:4,000 for IP samples), mouse HRP-anti-HA (BioLegend 901519, 1:1,000 dilution), HRP-anti-Myc monoclonal antibodies (BioLegend 626803, 1:2000 dilution), and HRP-Streptavidin (Jackson ImmunoResearch #016-030-084, 1:10,000 dilution). HRP-conjugated donkey anti-mouse IgG (Jackson ImmunoResearch #715-035-150, 1:10,000 dilution) was used for anti-tubulin and anti-GFP. HRP-conjugated goat anti-rabbit IgG (Thermo-Fisher #31462, 1:10,000 dilution) was used to detect anti-H2A and anti-H3. HRP-conjugated goat anti-rat IgG (Pierce #31470, 1:10,000 dilution) was used to detect anti-RGA (DU18). Chemiluminescent signals were detected and quantified by iBright FL1500 (Invitrogen).

The Phos-tag gel blot analysis was performed using 6% SDS-PAGE gel containing 25 μM Phos-tag Acrylamide reagent (FUJIFILM Wako Chemicals USA Corp. #AAL-107) and 50 mM MnCl_2_ as described^42^.

### In vitro pulldown and co-IP assays

In vitro pulldown assay was performed following the procedures published previously with slight modifications^24,55^. Recombinant proteins (GST, GST-BZR1, GST-PIF3, MBP and MBP-MED15^N^) expressed in BL21-CodonPlus (DE3)-RIL (Agilent Technologies) were purified with glutathione beads or amylose resin. GST or MBP-fusion proteins bound to glutathione or amylose beads were then used separately to pull down His-FLAG-RGA^GKG^ from protein extracts of transgenic Arabidopsis in specified genetic backgrounds.

For in vivo co-IP assays, protein complexes in total extracts from *Arabidopsis* or *N. benthamiana* were immunoprecipitated using anti-FLAG M2 Affinity Agarose Gel (MilliporeSigma, A2220), rabbit anti-Myc polyclonal antibody-conjugated Agarose Affinity Gel (A7470, MilliporeSigma) or GFP-Trap Magnetic Agarose beads (gtma-20; ChromoTek), as described previously^24,55^.

### TurboID-based proximity labeling and purification of biotinylated proteins

Identification of RGA proximity proteome was performed as described previously^79^ with some modifications. Ten-day-old Arabidopsis seedlings grown in liquid MS media were incubated with 100 μM biotin for 3 h before harvesting. One gram of finely ground tissue was resuspended in 2.5 mL extraction buffer (50 mM Tris pH 7.5, 150 mM NaCl, 0.1% SDS, 1% Triton-X-100, 0.5% Na-deoxycholate, 1 mM EDTA, 1 mM DTT, 1x SigmaFAST, 1 mM PMSF and 20 μM MG132). After centrifugation (15000g, 10 min) at 4°C, supernatants were transferred to fresh tubes. Free biotin was removed using PD-10 desalting columns (GE Healthcare G-25M, #17085101) and proteins were eluted with 3.5mL equilibration buffer (extraction buffer without SigmaFAST, PMSF or MG132). The eluted proteins (5 mg) were incubated with 140 μL Pierce™ Streptavidin Magnetic Bead slurry for 3h at 4°C with rotation. Beads were washed sequentially with 1 mL buffer as follows: 2x with extraction buffer, 2x with equilibration buffer, 1x with 1M KCl, and 2x with extraction buffer at 4°C. Three biological replicates of proteins on beads were subject to tryptic digestion as described^79^.

### LC-MS/MS and data analysis for RGA proximitome

Liquid chromatography and mass spec acquisition was configured as described for an Orbitrap Eclipse^79^ with the following modifications. LC-MS/MS acquisition carried out on a Orbitrap Eclipse Tribrid mass spectrometer (Thermo Fisher), equipped with an Easy LC 1200 UPLC liquid chromatography system (Thermo Fisher). Peptides were first trapped using a trapping column (Acclaim PepMap 100 C18 HPLC, 75 μm particle size, 2 cm bed length), then separated using analytical column AUR4-25075C18, 25CM Aurora Series Emitter Column (25 cm x 75 µm, 1.7 µm C18) (IonOpticks). The flow rate was 300 nL/min, and a 120-min gradient was used. Peptides were eluted by a gradient from 3 to 28 % solvent B (80 % acetonitrile, 0.1 % formic acid) over 106 min and from 28 to 44 % solvent B over 15 min, followed by a short wash (9 min) at 90 % solvent B.

LFQ analysis was carried out as described^79^. Briefly, raw data were processed using FragPipe (version 22.0). Using MSFragger (version 4.1), database searching was performed against the Araport11 *Arabidopsis thaliana* proteome (2/14/2025 version, 48,354 entries, https://www.arabidopsis.org/) plus TurboID protein sequence. IonQuant (version 1.10.27) was used for quantification, with MaxLFQ enabled, a minimum ion count of two, and match-between-runs enabled. LFQ intensities were imported to Perseus (version 2.1.3) where a Student’s t-test was executed to identify candidate proteins that are enriched by the RGA-TurboID in comparison to the GFP-TurboID control. The t-test settings were the following: ‘Permutation-based FDR’, ‘FDR=0.01’, S0=0.8, ‘Report q-value’, ‘Number of Randomizations=250’ and ‘-log10 p-value’. Volcano plot was generated by GraphPad Prism 8. GO term analysis was performed using Protein Analysis Through Evolutionary Relationships (PANTHER 19.0)^80^, and the plot was generated with SRplot^81^. RGA interaction network analysis was performed using Search Tool for the Retrieval of Interacting Genes/Proteins (STRING 12.0)^82^, followed by Cytoscape 3.10.4^83^.

### Protein purification and identification of PTM sites by online liquid chromatography tandem MS (MS/MS) analyses

His-FLAG-RGA^GKG^ was purified from *P_RGA_:His-FLAG-RGA^GKG^* transgenic *Arabidopsis* lines in the *ga1-13* and *ga1-13 cdk8-1* backgrounds by the tandem affinity purification procedures described previously^84^. Affinity-purified proteins were trypsin-digested, and peptides were analyzed by online LC-electrospray ionization (ESI) tandem MS [electron-transfer dissociation (ETD) and collisionally activated dissociation (CAD) MS/MS] using a Thermo™ Orbitrap Fusion Lumos™ Tribrid™ mass spectrometer equipped with ETD^85^. Full MS scans (300–2,000 m/z) were acquired using the Orbitrap (120k resolution) and linear ion trap mass analyzers. Peptide monoisotopic peak detection was enabled. Ions 300–1,500 m/z were selected for fragmentation using data-dependent acquisition and a three-second cycle. Low charge ions (z = 1 and 2) were selected for stepped HCD fragmentation (20, 25, 30% NCE) and Orbitrap MS/MS analysis at 7.5k resolution. Higher charge ions (z = 3 to 7) were selected twice, first for CAD fragmentation (35% NCE, 10 ms activation, q = 0.25) and linear ion trap MS/MS analysis, then for ETD fragmentation (using calibrated reaction times) and linear ion trap MS/MS analysis. Precursor ions were isolated by the quadrupole using a 3 m/z width. Dynamic exclusion was implemented as a repeat count of 2, repeat duration count of 10 s, and exclusion duration of 10 s (replicate 1), or as a repeat count of 3, repeat duration count of 16 s, and exclusion duration count of 16 s (replicates 2 and 3).

MSFragger (FragPipe v24.0)^86^ was used to search data against a database containing 6His-3xFLAG-RGA^GKG^ with decoys. Search settings included +/- 5 ppm precursor error tolerance, 20 ppm fragment mass tolerance, and fully specific trypsin digest allowing 2 missed cleavages. Alkylation of cysteine residues was a fixed modification. Variable modifications included methionine oxidation, phosphorylation of serine, threonine, and tyrosine, *O*-fucosylation of serine and threonine, *O*-GlcNAcylation of serine and threonine, and *O*-hexosylation of serine and threonine. Spectral matches were manually validated using MS/MS spectra, and modification site localization was confirmed by manual inspection of the spectra^87^. Peptides were quantified by integrating the peak areas for all detected charge states (including ^13^C isotopes) using Thermo Scientific Xcalibur™ Qual Browser (v4.2.47). PTM abundances were calculated relative to the total mass area of all modification states detected for the peptide, averaged across replicate samples, and rounded to the nearest tenth decimal^42^. PTMs were only considered if MS/MS data supported full or partial site-localization.

### Accession numbers

Sequence information for Arabidopsis genes included in this article can be found in the GenBank/EMBL data libraries under accession numbers *RGA* (AT2G01570), *CDK8* (AT5G63610), *MED15A* (also known as *NON-RECOGNITION-OF-BTH 4*, AT1G15780.1), *MED12* (AT4G00450), *MED13* (AT1G55325), *SEUSS* (AT1G43850), *LEUNIG* (AT4G32551), *SCL3* (AT1G50420), *GID1B* (AT3G63010), *EXP8* (AT2G40610), *IQD22* (AT4G23060), *Exp-PT1* (AT2G45900), *GRF8* (AT4G24150), *XERICO* (AT2G04240), *BOI* (AT4G19700), *PMEI13* (AT4G15750), *IAA16* (AT3G04730), *GH3.3* (AT2G23170), *FLA9* (AT1G03870), *PP2A* (AT1G13320), *H2A* (AT1G51060), *BZR1* (AT1G75080) and *PIF3* (AT1G09530).

## RESOURCE AVAILABILITY

### Lead contact

Requests for further information and resources should be directed to, and will be fulfilled by, the lead contact, Tai-ping Sun.

### Materials availability

Materials generated in this study are available from the lead contact upon request. Material that can be shared will be released via a material transfer agreement.

### Data availability

The mass spectrometry proteomics data have been deposited to the ProteomeXchange Consortium via the PRIDE^88^ partner repository with the dataset identifier PXD077697 for RGA PTMs (Project DOI: 10.6019/PXD077697), and PXD078134 for RGA proximitome.

## Acknowledgements

We thank Dr. Clint Chapple for providing the *cdk8* mutants and CDK8/CDK8^D176A^ constructs, and Dr. Liang Jiang for providing the plasmid 3xHA-TurboID-NLS_pCDNA3. We thank Dr. Mark Ross for help and guidance investigating the MS/MS data. This work was supported by the National Institutes of Health (GM100051 and GM150029 to TPS, GM037537 to DFH, and S10OD030441 to SLX), the National Science Foundation (MCB-2416564 to TPS), the Carnegie Endowment Fund to the Carnegie Mass Spectrometry Facility, and the Hargitt Fellowship to XH.

## Author Contributions

T-pS and XH conceived and designed the research project. XH performed most molecular biology, genetics and biochemical analyses, HC participated in Y2H analysis, and XH, HC and T-pS analyzed the data and generated figures. LR, JS and DFH performed and analyzed the MS/MS data for RGA PTMs. AVR and SLX performed and processed the MS/MS data for RGA-TbID samples. T-pS and XH wrote the manuscript with input from all co-authors.

