## Supplemental Figures for "CDK8 phosphorylation of DELLA limits Mediator recruitment in gibberellin signaling"

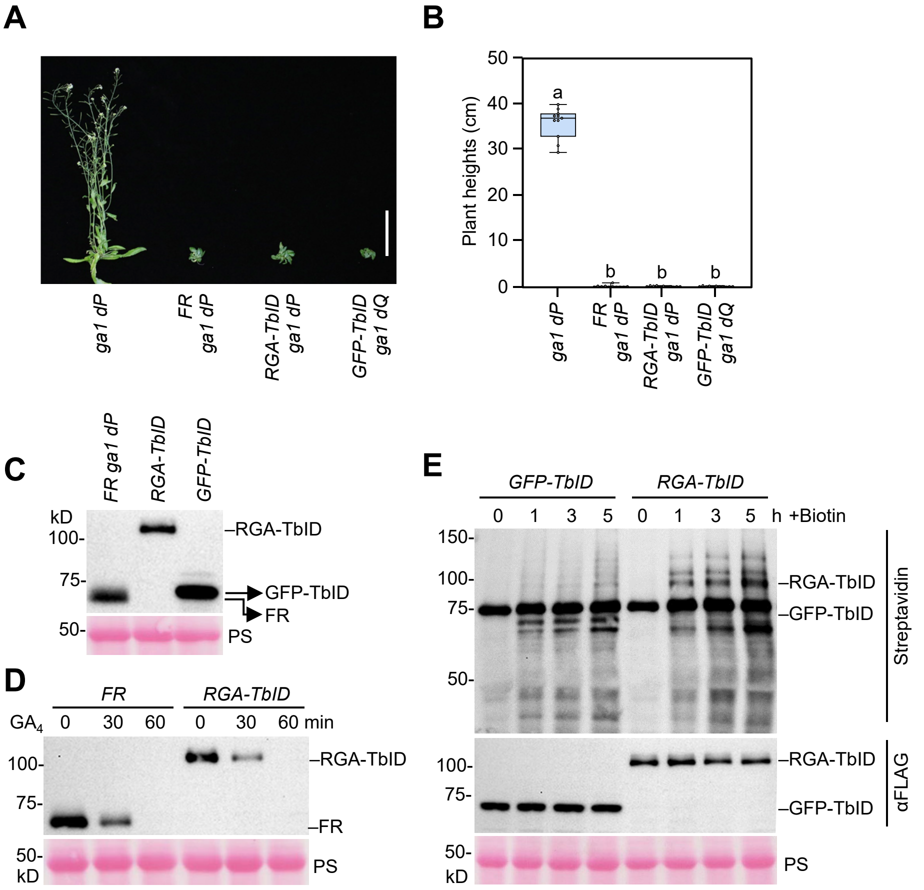


**Figure S1. TbID-based labeling of RGA proximity proteome in *Arabidopsis*.** (A)-(C) RGA-TbID is functional in planta. (A) $P_{RGA}$*:His-FLAG-RGA-TbID* in *ga1 dP* background (named *RGA-TbID* line, #1) was functionally like $P_{RGA}$*:His-FLAG*-RGA (line #5) to restore the dwarf phenotype. $P_{RGA}$*:His-FLAG-NLS* (nuclear localization sequence)-*GFP-TbID* transgenic line in the *ga1 dQ* (quadruple *della* with WT *RGA*) background (named *GFP-TbID*, line #13) was included as a control. Photo of representative 49d-old plants was taken. Bar = 10 cm. (B) Plant heights are shown in the boxplot. n=11. Center lines and box edges are medians and the lower/upper quartiles, respectively. Whiskers extend to the lowest and highest data points within 1.5× interquartile range (IQR) below and above the lower and upper quartiles, respectively. Different letters above the bars represent significant differences ($p < 0.01$) as determined by two-tailed Tukey's HSD mean separation test. The phenotypic analysis was repeated two times with similar results. (C) Immunoblot analysis using an anti-FLAG antibody showing that FLAG-RGA, RGA-TbID and GFP-TbID are expressed at similar levels. (D) RGA-TbID was degraded similarly as FLAG-RGA after treatment with 0.5 µM $GA_4$. (E) Immunoblot analysis showing biotinylated proteins in RGA-TbID and GFP-TbID lines after incubation with 100 µM biotin for 0h-5h as labeled. The blots were probed with HRP-Streptavidin or anti-FLAG antibody. In (C-E), Ponceau S (PS)-stained blots showing equal loading.

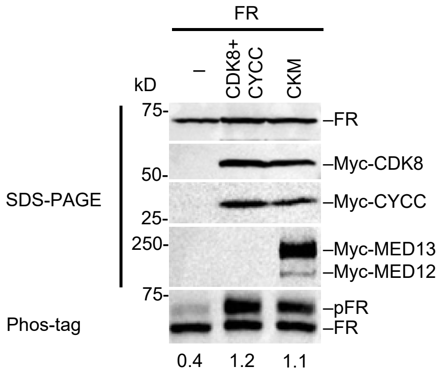


**Figure S2. Co-expression of MED12 and MED13 did not further increase RGA phosphorylation by CDK8 and CYCC in *N. benthamiana*.** FLAG-RGA (FR) was expressed alone (–) or together with Myc-CDK8 and -CYCC or all four subunits of CDK8 Kinase module (CKM, including CDK8, CYCC, MED12 and MED13) in *N. benthamiana*. The ratios of phosphorylated FR (pFR)/unphosphorylated FR are shown below the Phos-tag gel blot.

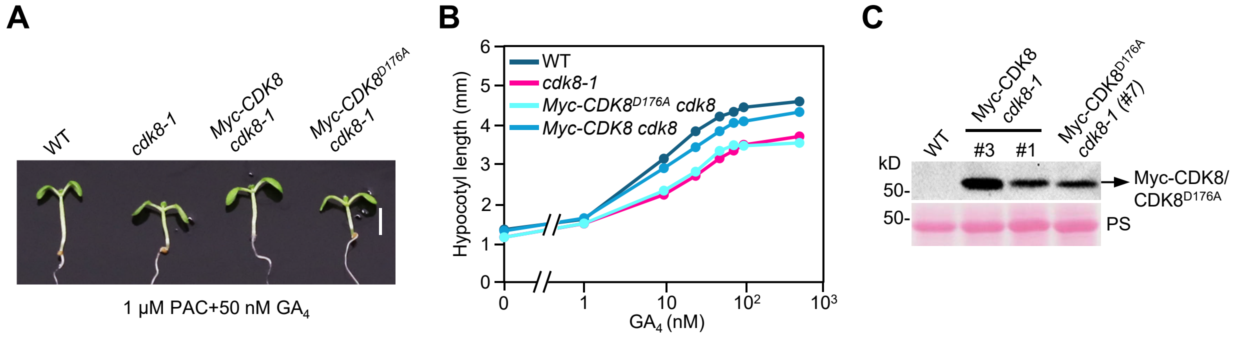


**Figure S3. Phenotypes of *cdk8* mutants and transgenic lines.** (A)-(B) *35S:Myc-CDK8*, but not *35S:Myc-CDK8$^{D176A}$*, rescued the GA response of *cdk8-1*. Seedlings were grown in medium containing 1 μM PAC and varying concentrations of $GA_4$. The photo was taken and hypocotyl lengths were measured at day 9. (A) Bar = 3 mm. (B) Average hypocotyl lengths. Means ± SE. n=12. (C) Protein blot showing similar levels of Myc-CDK8 and Myc-CDK8$^{D176A}$ in the *Myc-CDK8 cdk8-1* transgenic line (#1) and *Myc-CDK8$^{D176A}$ cdk8-1* line (#7), which were used in (A-B). Another *Myc-CDK8 cdk8-1* transgenic line (#3) that accumulates higher amounts of Myc-CDK8 was used in **Figure 4**.

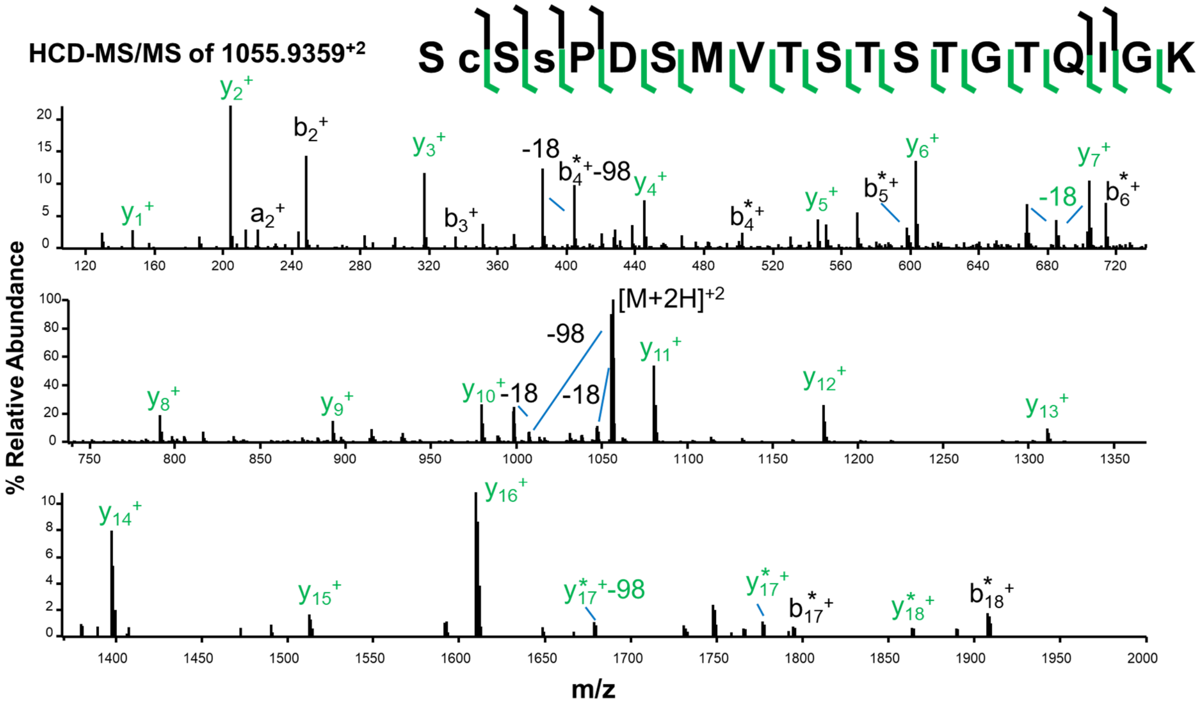


**Figure S4. High-resolution HCD-MS/MS of fully tryptic Poly-S/T region identifying pS170.** Example from *Arabidopsis ga1-13* replicate #1. Lowercase "c" denotes alkylated cysteine, lowercase "s" denotes phosphorylated serine, * denotes +80 Da-shifted (phosphorylated) ions, -98 denotes loss of phosphoric acid, -18 denotes water loss.

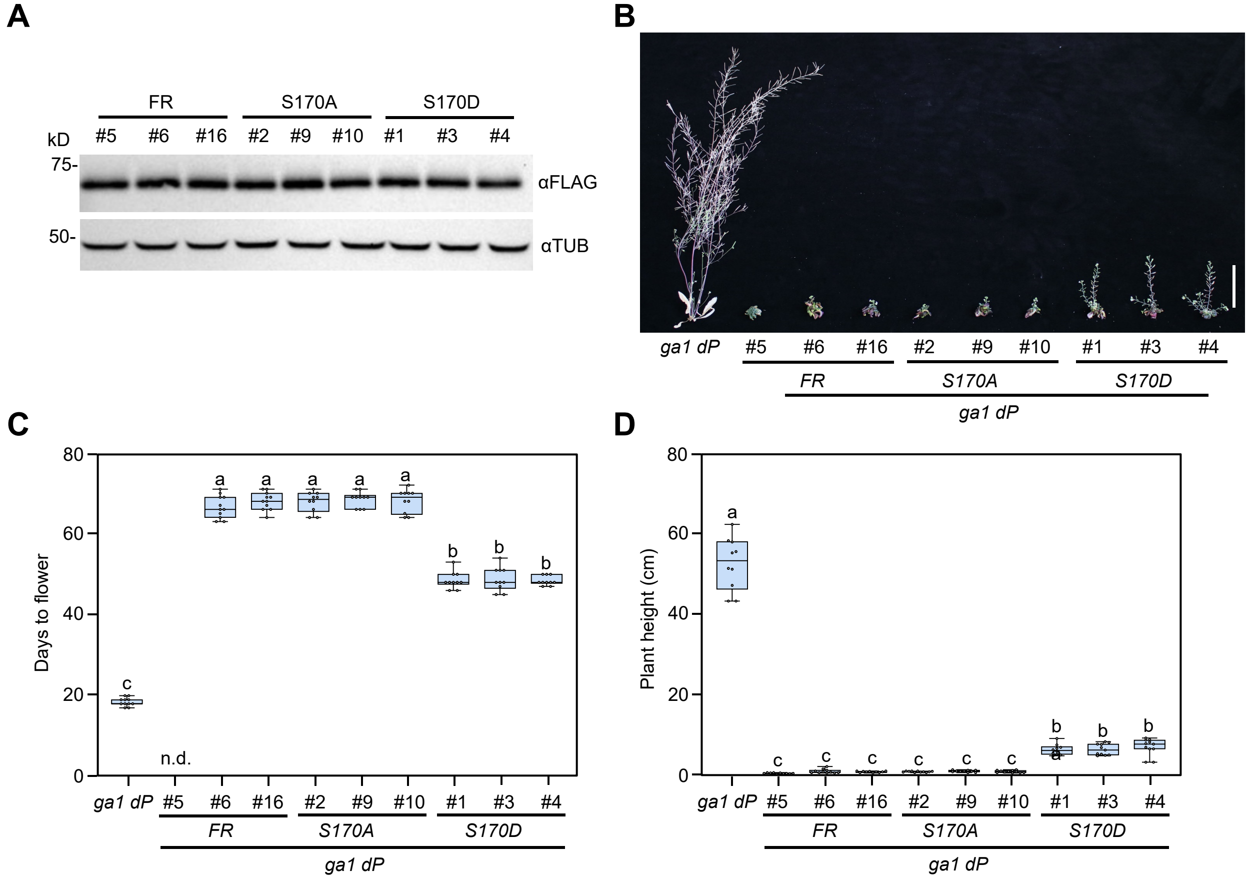


**Figure S5. S170D caused reduced RGA activity in planta.** (A) All transgenic lines accumulated similar levels of His-FLAG-RGA/rga$^{S170A}$/rga$^{S170D}$. Immunoblots containing total proteins from $P_{RGA}$*:His-FLAG-RGA/rga ga1 dP* lines were probed with anti-FLAG and anti-TUB antibodies, separately. (B)-(D) Phenotypes of $P_{RGA}$*:His-FLAG-RGA/rga* transgenic lines in the *ga1 dP* background. (B) Representative 93d-old plants under LD. Bar = 5 cm. (C-D) Boxplots showing flowering time and plant heights of different lines as labeled. n=10-11. Center lines and box edges are medians and the lower/upper quartiles, respectively. Whiskers extend to the lowest and highest data points within 1.5× interquartile range (IQR) below and above the lower and upper quartiles, respectively. Different letters above the bars represent significant differences ($p < 0.01$) as determined by two-tailed Tukey's HSD test. The phenotypic analysis was repeated two times with similar results.

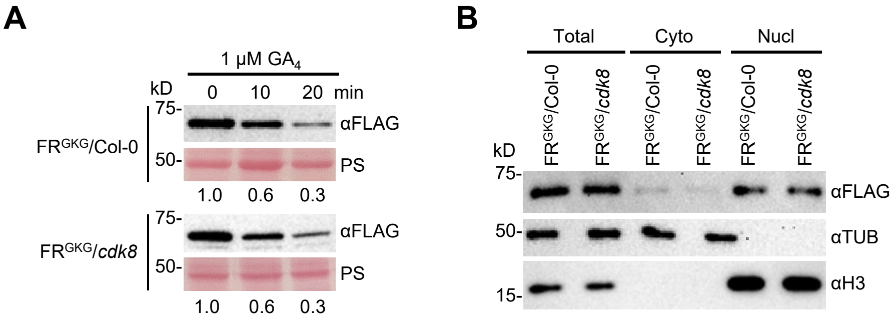


**Figure S6. CDK8 mutation did not alter GA-induced degradation or nuclear localization of RGA in *Arabidopsis*.** (A) *cdk8-1* did not alter GA-induced degradation of RGA in *Arabidopsis*. Protein blots containing protein extracted from 10d-old transgenic *Arabidopsis* $P_{RGA}$*:His-FLAG-RGA*$^{GKG}$ in Col-0 or *cdk8-1* background that were grown in the presence of 1 μM PAC, and then were treated with 1 μM $GA_4$ at indicated time points. The blots were probed with an anti-FLAG antibody. Ponceau S (PS)-stained blot indicated similar sample loading. (B) *cdk8-1* did not affect nuclear localization. Immunoblots containing total and fractionated proteins from transgenic *Arabidopsis* $P_{RGA}$*:His-FLAG-RGA*$^{GKG}$ in Col-0 or *cdk8-1* background grown in the presence of 1 μM PAC for 10 days were probed with anti-FLAG, anti-histone H3 and anti-tubulin (TUB) antibodies, separately. In (A-B), the assay was repeated two times with similar results.

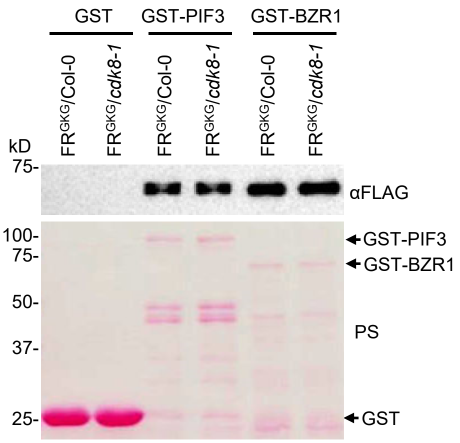


**Figure S7. In vitro pulldown assays to examine TF interaction with FLAG-RGA in WT and *cdk8-1* backgrounds.** The images of Ponceau S-stained blots show the GST and GST-BZR1, GST-PIF3 proteins used in the pull-down assays in **Figure 7A**. The assay was repeated two times with similar results.

| Sample # | pGBKT7 (Bait) | pGADT7 (Prey) | pH3-Myc |
|---|---|---|---|
| 1 | – | – | CDK8$^{D176A}$/CYCC |
| 2 | – | – | CDK8 /CYCC |
| 3 | – | RGA | CDK8$^{D176A}$/CYCC |
| 4 | – | RGA | CDK8 /CYCC |
| 5 | MED15 | – | CDK8$^{D176A}$/CYCC |
| 6 | MED15 | – | CDK8 /CYCC |
| 7 | MED15 | RGA | – |
| 8 | MED15 | RGA | CDK8$^{D176A}$/CYCC |
| 9 | MED15 | RGA | CDK8 /CYCC |

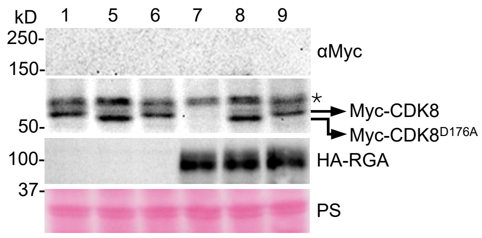


**Figure S8. CKM weakened RGA-MED15 interaction.** Immunoblot analyses showing similar levels of RGA, CDK8 and CDK8$^{D176A}$ among different samples in the Y3H assays in **Figure 8A**. MED15 was not detectable. Protein blots containing extracts from yeast cells were probed with anti-Myc and anti-HA antibodies, separately. Ponceau S (PS)-stained blots showing similar loading. *, non-specific background band. Two biological repeats showed similar results.

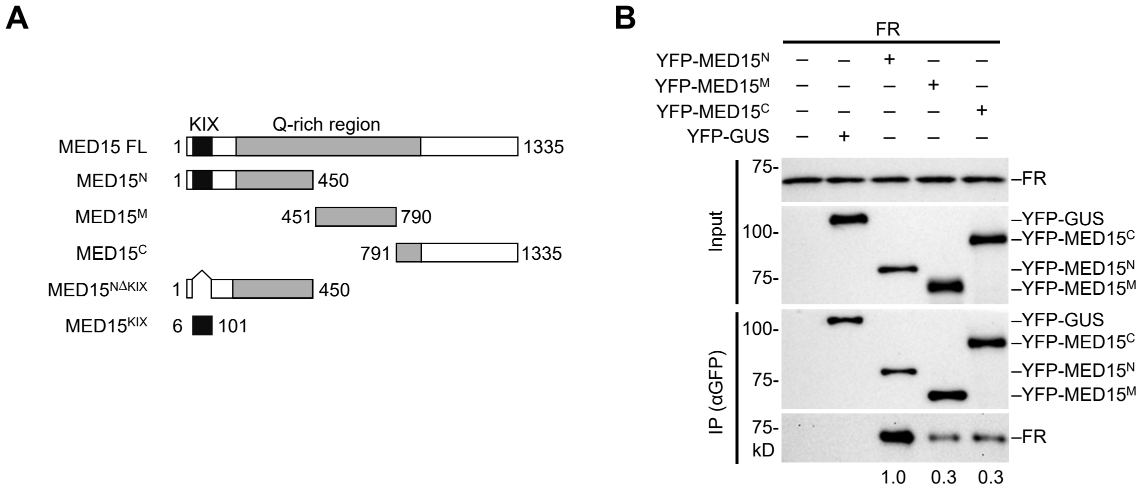


**Figure S9. Mapping RGA interaction domain in MED15 by co-IP assays.** (A) Schematics of full-length (FL) and truncated MED15. (B) Co-IP assays showed that all three MED15 fragments (N, M and C) bind to RGA, although MED15$^{N}$ displays the highest affinity. FLAG-RGA was expressed alone (–) or together with YFP-MED15$^{N}$, MED15$^{M}$ or MED15$^{C}$ in *N. benthamiana*. YFP-GUS was included as a negative control. YFP-MED15 fragments or -GUS were immunoprecipitated using anti-GFP agarose. Top panel (Input): Immunoblots containing total protein extracts. Bottom panel (αGFP immunoprecipitated samples): Immunoblots containing IP eluates were probed with anti-FLAG or anti-GFP antibodies. Representative images of 2 biological repeats are shown.

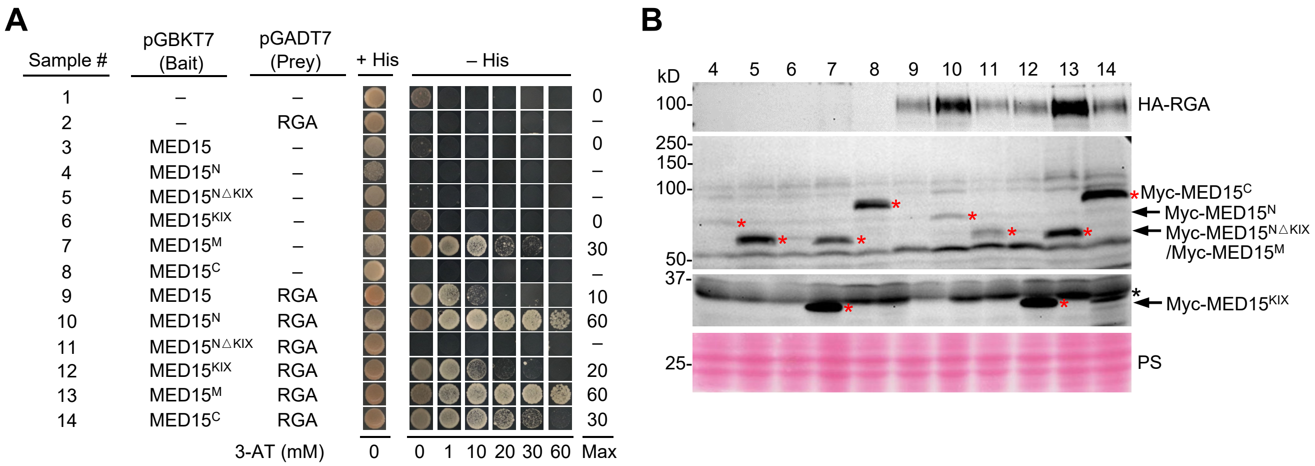


**Figure S10. Mapping RGA interaction domain in MED15 by Y2H assays.** (A) Y2H assays showing RGA interaction with all three MED15 fragments (N, M and C) and the KIX domain. MED15-FL and truncated fragments were fused to the Gal4 DB domain as the baits, and the full-length RGA was fused to Gal4 AD domain as the prey. Interaction of DB and AD fusion proteins in the PJ69-4A yeast cells was scored by the relative growth in –His media containing varying concentrations of 3-AT as labeled. – indicates empty vector, or no growth at 0 mM 3-AT. Max, detectable cell growth at maximum 3-AT concentration. (B), Immunoblot analyses showing relative levels of RGA, and different MED15 fragments (indicated by the red asterisks) across samples. Protein blots containing extracts from yeast cells were probed with anti-Myc and anti-HA antibodies, separately. Ponceau S (PS)-stained blots showing similar loading. *, non-specific background band. Two biological repeats showed similar results.

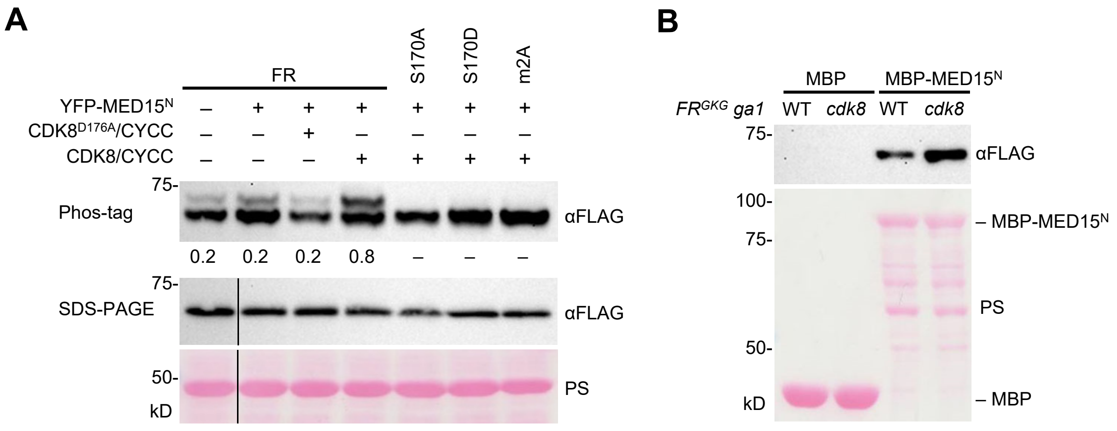


**Figure S11. Input samples for co-IP and pulldown assays.** (A) Phos-tag gel blot showing elevated levels of phosphorylated FLAG-RGA when CDK8 and CYCC were co-expressed in *N. bethamiana*. The blot containing the input samples used in the co-IP assays in **Figure 9**. (B) In vitro pulldown assays to examine $MED15^{N}$ interaction with $FLAG\text{-}RGA^{GKG}$ *ga1* in WT and *cdk8-1* background. The images of Ponceau S-stained blots show the MBP and $MBP\text{-}MED15^{N}$ proteins used in the pull-down assays in **Figure 8B**. The assay was repeated two times with similar results.
