## Supplementary material for "CDK8 phosphorylation of DELLA limits Mediator recruitment in gibberellin signaling": Table S3

### Table S3. Posttranslational modifications in His-FLAG-RGA$^{GKG}$ from *ga1* and *ga1 cdk8*

| Residues [a] | Peptide Sequence [b] | PTM | Sites [c] | Relative Abundance (%) | |
|---|---|---|---|---|---|
| | | | | ***ga1-13*** | ***ga1-13 cdk8-1*** |
| 12-26(28) | LSNHGTSs$^{Phos}$SSSSISK(DK) | 1 Phos | S19 | 5.0 | 4.9 |
| | LSNHGTSSs$^{Phos}$SSSISK(DK) | 1 Phos | S20 | | |
| | LSNHGTSSSs$^{Phos}$SSISK(DK) | 1 Phos | S21 | | |
| | LSNHG[TSSSSSSIS]$^{2Phos}$K(DK) | 2 Phos | Not mapped to a specific residue | | |
| | LSNHGt$^{GlcNAc}$SSSSSSISK(DK) | 1 GlcNAc | T17 | 2.5 | 2.2 |
| | LSNHGTs$^{GlcNAc}$SSSSSISK(DK) | 1 GlcNAc | S18 | | |
| | LSNHGTSs$^{GlcNAc}$SSSSISK(DK) | 1 GlcNAc | S19 | | |
| (165)167-185 | (LK)SCSs$^{Phos}$PDSMVTSTSTGTQIGK | 1 Phos | S170 | 32.9 | 16.0 |
| | (LK)[SCSSPDSMVTSTSTGT]$^{GlcNAc}$QIGK | 1 GlcNAc | Not mapped to a specific residue | 7.9 | 7.9 |
| 186-207 | GVIGTTVTTTTTt$^{GlcNAc}$TTAAGESTR | 1 GlcNAc | T198 | 69.4 | 67.4 |
| | GVIGTTVTTTTTTt$^{GlcNAc}$TAAGESTR | 1 GlcNAc | T199 | | |
| | GVIGTTVTTTTTTTt$^{GlcNAc}$AAGESTR | 1 GlcNAc | T200 | | |
| | GVIG[TTVTTTTTTTTAAGEST]$^{2GlcNAc}$R | 2 GlcNAc | Not mapped to a specific residue | | |
| | GVIG[TTVTTTTTTTTAAGEST]$^{Hex}$R | 1 Hex | Not mapped to a specific residue | 8.6 | 10.7 |

[a] Residues in WT RGA, with numbers in parentheses indicating the residues of longer peptides resulting from missed trypsin cleavage, and abundances of fully and partially tryptic peptides were combined. [b] Brackets indicate the region in the peptide that contains the PTM when it could not be mapped to a specific amino acid. [c] Sites were determined by signature CAD and/or ETD fragmentation, including diagnostic ions. Peptide abundances were determined from MS1 peak areas and PTM levels are reported as the % of the total peptide abundance, including all modified forms detected. Total abundances were calculated as [(modified peptide MS1 peak area)/(sum of all modified peptides + unmodified peptide peak areas)] x 100. The percentage values are the average of three biological replicates.
